# When mice think twice: optimal control explains when and how mice overcome default biases

**DOI:** 10.1101/2023.01.04.522806

**Authors:** K. Barc, C. Schreiweis, M. Euvrard, J. Mattout, E. Burguière, J. Daunizeau

**Affiliations:** Paris Brain Institute - ICM, Hôpital de la Pitié Salpêtrière, Paris, France; Lyon Neuroscience Research Center, Univ Claude Bernard Lyon 1, Inserm, CNRS, UMR 5292 / U-1028, Lyon, France; ETH, Zurich, Switzerland

## Abstract

Decisions permeate every aspect of our lives but decision-making policies seem to vary tremendously across time and individuals. Rather than processing all decision-relevant information, we sometimes rely on habitual and/or intuitive responses, which can lead to irrational biases and errors. This raises the twofold issue of whether a single normative principle can (i) determine when cognitive control is engaged, and (ii) account for the diverse behavioral signature of cognitive control regulation. In line with recent theoretical proposals, our working hypothesis is that cognitive control is flexibly deployed to optimize a cost-benefit trade-off. Using Markov Decision Processes, we derive the optimal online control allocation policy in standard decision-making tasks, under arbitrary default biases. This bridges the gap between motivational theories of cognitive control regulation and accumulation-to-bound models of decision making. Under the model, response times, default response rates, decision accuracy and interference effects are not independent behavioral phenomena, but distinct manifestations of the same underlying arbitration process. We then test the distinct behavioral predictions of the model using a customized pseudo-ecological set-up, whereby mice engage with a self-administered perceptual decision-making task over several weeks. In this context, processing decision-relevant cues engages cognitive control, whereas default responses are determined by learned, cue-independent, action-outcome associations. The model accurately predicts both performance and response time interference effects, their interactions, as well as their modulation by reward and task demands. This work provides novel computational means for assessing the brain circuits that operate the arbitration between default and controlled decision processes, in rodent models of both healthy and neuropsychiatric conditions.

## INTRODUCTION

Decisions permeate every aspect of our lives - what to eat, where to live, etc. - but the way we make decisions varies tremendously. Rather than processing all decision-relevant information, we often rely on habitual and/or intuitive responses, which can lead to irrational biases and errors *(Kahneman et al., 1982; Schneider & Shiffrin, 1977)*. For example, a distracted driver may automatically follow the familiar route home despite intending to stop elsewhere, allowing a well-learned habit to override the current goal. Yet, we do not always follow the lead of automatic decision processes. So what mechanism regulates the use of default responses?

Heedless decisions can be triggered by psychobiological determinants such as stress *(Porcelli et al., 2012; Porcelli & Delgado, 2009)*, emotions *(Harlé & Sanfey, 2007; Martino et al., 2006; Sokol-Hessner et al., 2013)* or fatigue *(Blain et al., 2016; Wiehler et al., 2022)*. However, they also occur in the absence of such contextual factors, suggesting that they cannot be explained solely by transient fluctuations of physiological or affective states. On the one hand, standard accumulation-to-bound models of decision-making rather emphasize the role of unreliable or stochastic neural representations of decision-relevant information *(Drugowitsch et al., 2016; Wang & Busemeyer, 2016; Wyart & Koechlin, 2016)*. However, they cannot, by themselves, explain the brain’s flexible strategy for suppressing or relying on default responses. What is lacking here are higher-level cognitive computations that arbitrate conflicts between distinct behavioral processes under changing goals and constraints. On the other hand, opponent-process theories suggest that spontaneous automatic responses reflect an insufficient engagement of *cognitive control* - the effortful process of monitoring and adjusting ongoing information processing according to current goals *(Badre, 2025; Li et al., 2017; McGuire & Botvinick, 2010; Wu et al., 2016)*. In humans, early empirical demonstrations of this idea came from variants of the Stroop task, in which a prepotent or default response (e.g., reading a word) interferes with the task-relevant or instrumental response (e.g., naming its ink color). Here, engaging cognitive control enables slow but accurate processing of the task-relevant information, which tends to override the default response *(Cohen et al., 1990; Frank, 2006; Frank et al., 2007; Kalanthroff et al., 2018)*. However, the issue of how cognitive control is itself regulated remains open. A recent theoretical proposal is that cognitive control is deployed to the extent that its expected benefit outweighs its prospective costs *(Kool et al., 2010; Lieder et al., 2018; Shenhav et al., 2013a)*. This strategic deployment of cognitive control explains the contextual modulation of Stroop-like interferences effects by reward and task demands *(Egner & Hirsch, 2005; Krebs et al., 2010; Veling & Aarts, 2010)*. A promising next step is to complement this motivational framework with optimal accumulation-to-bound models, which formalize how decision thresholds should be adjusted to balance speed and accuracy *(Bénon et al., 2024; Drugowitsch et al., 2012, 2019; Tajima et al., 2016)*. This would explain how the broad range of behavioral signatures of cognitive control regulation – including response times, default response rates and decision accuracy – may emerge from the same cost-benefit arbitration process. However, this theoretical synthesis has not been performed yet. And whether a single computational mechanism can simultaneously account for these diverse phenomena has never been tested experimentally.

In principle, computational models of cognitive control regulation apply to, e.g., primates or rodents. In combination with, for example, targeted chemogenetic or optogenetic manipulations of decision-related subsystems in the brain, they would enable quantitative computational decompositions of overt behavior with potential relevance for translational neuropsychiatric approaches *(Johnson et al., 2013; Robble et al., 2021)*. Of course, a critical issue here is whether a given animal species is equipped with the cognitive apparatus that is necessary to flexibly deploy cognitive control. In non-human primates, there is clear evidence that various monkey species can strategically mobilize cognitive control *(Brown & Hampton, 2020; De Petrillo et al., 2022)*. In comparison, evidence for rodents’ ability to flexibly deploy cognitive control is still debated, at least in mice *(Evans et al., 2019; Park et al., 2025)*. For example, even after extensive training on simple perceptual decision-making tasks, mice exhibit unequal performances from trial to trial, occasionally returning to suboptimal default (e.g., perseverative) responses *(Ashwood et al., 2022; Roy et al., 2021; Urai, 2026)*. Nevertheless, this within-individual behavioral variability may in fact suggest that mice can, under some circumstances, override their default bias. However, the mechanisms governing when and how this occurs remain unknown. Moreover, it is unclear whether a single computational mechanism can simultaneously account for the multiple behavioral consequences of control allocation.

In this context, the contribution of this work is twofold. First, we extend previous computational work on the motivational regulation of cognitive control *(Bénon et al., 2024; Griffiths et al., 2015; Lee & Daunizeau, 2021; Lieder et al., 2018; Shenhav et al., 2013a)*, which essentially frames the allocation of cognitive control as a cost/benefit optimization problem. Here, we rely on Markov decision processes to derive the optimal control allocation policy under possibly conflicting default and controlled responses, and as a function of reward at stake and task difficulty. Under the model, response times, default response rates, decision accuracy and interference effects are not independent behavioral phenomena, but distinct manifestations of the same underlying arbitration process. Second, we test whether the model accurately captures default-induced interference effects in both performance and response time, their interaction, as well as their modulation by motivational factors. To do this, we developed a self-administered, automated, operant conditioning paradigm. During several weeks, mice stay in a customized conditioning box, in which they obtain food only when they successfully engage with a deterministic, binary, visual decision task. In this context, processing the visual cue engages cognitive control, whereas default responses are determined by learned, cue-independent, action-outcome associations. Critically, we systematically varied decision difficulty (i.e., visual cue discriminability) and reward at stake (i.e., amount of food) and quantified the ensuing variations in default-induced interference effects. This enabled us to track, over multiple days and in a pseudo-ecological environment, how mice arbitrate between automatic and controlled responses, beyond variations in endogenous psycho-biological states (such as circadian cycle phase, hunger state, stress, etc.) that may otherwise confound decision mechanisms.

## MATERIALS AND METHODS

### Animals

All experimental procedures followed national and European guidelines and were approved by the institutional review boards (French Ministry of Higher Education, Research and Innovation; APAFiS Protocols no. 1418-2015120217347265 and 2021042017105235). Animals were bred and group-housed in the animal facilities of the Paris Brain Institute in Tecniplast ventilated polycarbonate cages under positive pressure with hard-wood bedding in groups of up to six animals per cage with *ad libitum* food and water access. The temperature was maintained at 21–23 °C and the relative humidity at 55 ± 10\% with a 12-h light/dark cycle (lights on/off at 8am and 8pm, respectively). The animals were maintained in a 12-hour light/dark cycle (lights on/off at 8:00am/8:00 pm, respectively), and had *ad libitum* food and water access. In this study, n = 25 adult male mice were tested.

### Experimental setup

Operant conditioning was conducted in custom-modified experimental chambers (ENV-007CTX, Med Associates, Vermont, USA) as previously described *(Benzina et al., 2021)*, in which each of the tested animals lives and performs the conditioning task 24/7 in an automated, self-initiated, self-paced manner. Seven such experimental operant chambers were operated in parallel in the same experimental room. Briefly, the rear wall of each chamber housed the feeder compartment (equipped with an infrared light beam to detect feeder entry crossing) and drinking water bottle holder, the front wall held two tactile screens, and a custom-adapted, homemade pair of black Plexiglas gates equipped with a pair of infrared (IR) beams. These beams allowed for tracking the locomotion of the mouse into or out of the area with the tactile screens. Precision reward pellets (20 mg LabTabTM AIN-76A rodent precision pellets, TestDiet, Richmond, USA) served as the sole nutrition during the entire experiment and all rewards had to be earned through successful performance of the task.

### Pre-training phases

Once placed in the operant conditioning chambers, the mice start with a pre-training phase of five different stages. In the first stage, a total of five precision pellets drop automatically once every minute if the previous pellet had successfully been recovered. In the second stage, the mouse is required to nose poke into the feeder compartment to trigger individual pellet delivery once per minute until a total of ten pellets has been received. In the third stage, crossing of the photobeams prompts the illumination of both tactile screens; the mouse is required to touch one of them to trigger pellet delivery in the feeder. There was no time limit for recovering the pellets at this stage. In the fourth stage, mice have to nose poke into the feeder in less than 15 seconds after screen touch in order to trigger pellet delivery. Having successfully recovered at least 66% of the pellets, mice passed into the last (fifth) pre-training stage. This consisted in a deterministic, binary, visually-cued, decision task. At each trial, both touch screens displayed a visual cue composed of ten white lines, either vertical or horizontal. The line orientation was randomized over trials (with an overall 50% frequency). To earn a precision pellet, mice need to associate each of the two visual cues with one screen touch, e.g., horizontal/vertical lines associated with the action of left/right screen touch, respectively. The cue was shown simultaneously on both touchscreens for two reasons: first, even if mice would only partially sample from one screen, they would have all information at their disposal to successfully solve the task; and, second, the mouse signals its decision (which screen it touches) independently from the way we present the visual cue. The cue-outcome contingencies were randomized across individuals and operant chambers. All trials were self-initiated and onsets of the visual cue on the tactile screens were triggered by the mouse crossing the photobeam-equipped gates. Mice were allowed a maximum of 60 seconds after cue onset to respond with a screen touch. Upon correct response, the mouse had 15 seconds to nose poke in the food dispenser in the rear wall for obtaining a food reward. In case of incorrect screen touch, mice were exposed to an aversive white light for five seconds. Additionally, following an incorrect response, the rewarded side was set opposite to the previous response side (so-called “corrective trials”). This design feature was primarily introduced to minimize mice’s natural response lateralization bias and hence facilitate learning of the task structure. In addition, this action-outcome contingency provided direct reinforcement for cue-independent lose-switch responses, thereby also indirectly promoting win-stay responses (in non-corrective trials). Note that cue and reward stay yoked throughout, i.e., following the cue in corrective trials still pays off. Past 60 seconds in the chamber compartment with the screens, the screens would switch off again; in these “time out” trials, mice then had to cross out and back into the feeder compartment to trigger the next trial. Trials where mice crossed in and out of the screen compartment within 60 seconds and without screen touch, were labelled as “aborted trials”.

### Main decision task

Having reached 66% correct responses over the past 40 trials of the fifth pre-training phase, mice transitioned into the main (decision making) task, where we systematically modulated decision difficulty and reward at stake. Difficulty levels consisted of variations of the visual cue, whereby the dominant line orientation (indicating the correct response) is rendered less salient. More precisely, we used different ratios of intersecting vertical and horizontal lines: the easiest difficulty level consisted of the same 10 vertical or 10 horizontal lines as in the fifth pre-training stage; the next difficulty level consisted of 9:1 intersecting vertical and horizontal lines, etc., until 5:5 intersecting horizontal and vertical lines, representing the highest and unsolvable difficulty level. Except for the control 5:5 difficulty level (which was rewarded randomly 50% on each side), the cue-outcome contingency was identical to that of the fifth pre-training stage. Upon a correct response, the mouse earned either one or three reward pellets (hereafter: low and high reward conditions, respectively). The reward condition was blocked into series of 60 consecutive trials, with 10 trials per difficulty level. Each trial began with a light indicating the current reward condition. Note that, as in the fifth pre-training phase, following an incorrect response, the rewarded side was set opposite to the animal’s previous choice (corrective trials).

Figure 1 below shows the operant chamber and associated task design, as well as summary statistics of mice behavior during the last training phase and the main experimental phase.

**Fig. 1.**
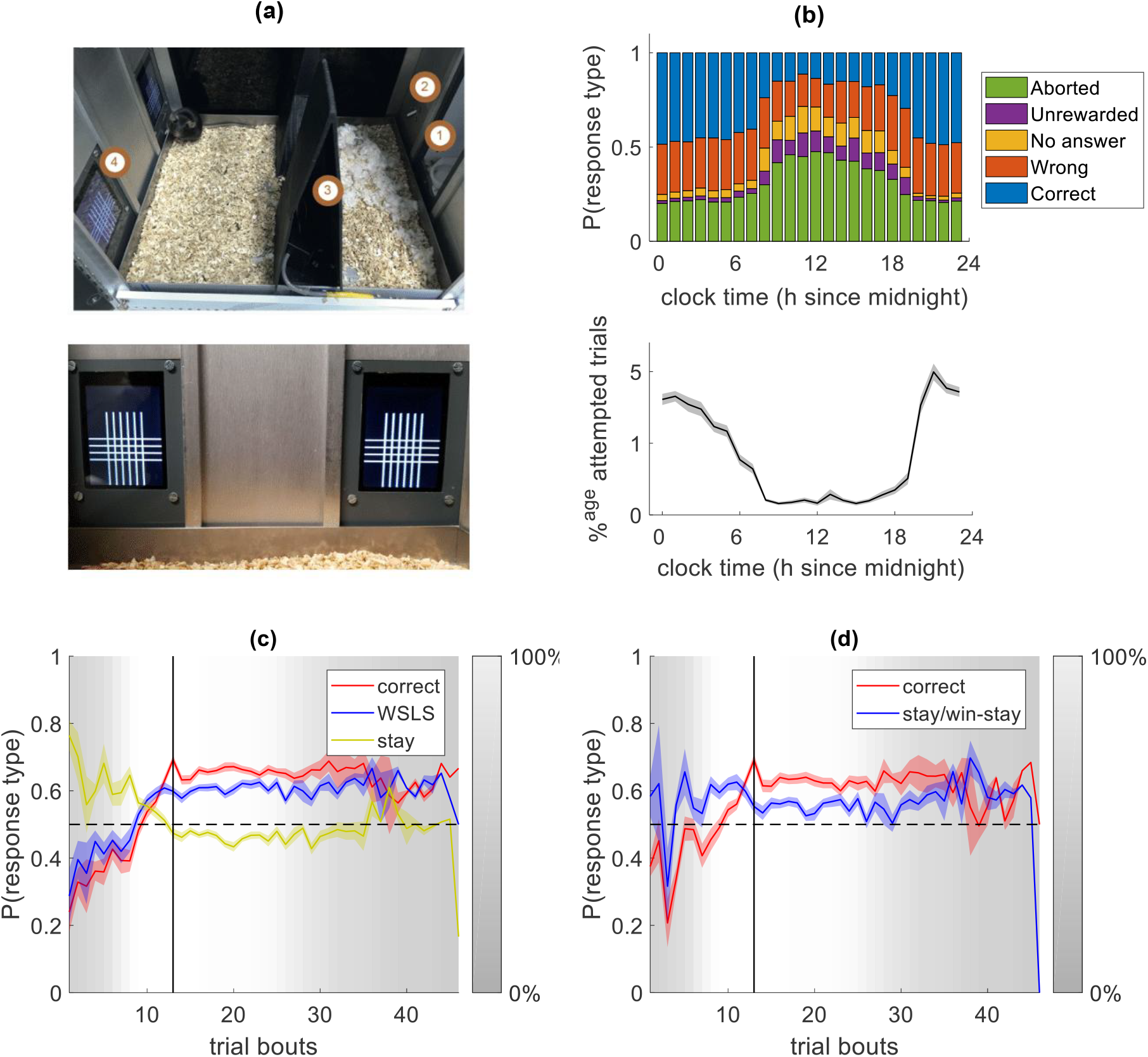
Experimental design and response rates. **(a)** Task apparatus. *Top:* Schematic of the experimental setup. (1) Food pellet dispenser; (2) lever used to obtain the reward; (3) entrance to the task zone, monitored by infrared detectors; and (4) tactile screens located within the task zone. *Bottom:* Close-up of the task zone showing the two symmetrical tactile screens positioned on the left and right sides. **(b)** *Top*: rate of response types across the circadian cycle. *Bottom*: percentage of attempted trials per clock hour. **(c)** Dynamics of correct (red), win-stay/lose-switch (blue) and perseverative/stay (yellow) response rates in valid trials across the final pre-training phase and the main experimental phase (each bout contains 200 valid trials). Shaded areas depict standard error around the mean (s.e.m.). Trial bouts are coregistered w.r.t. the onset of the main experimental phase (thick vertical black line). The lightness of the background reports the proportion of animals contributing to data within each bout, reflecting variability across mice in the duration of training and in the total number of trials completed during the main experimental phase. **(d)** Dynamics of correct (red), and perseverative/win-stay (blue) response rates in non-corrective trials across the final pre-training phase and the main experimental phase.

As in any ecological setting, behavior is determined by multiple imperatives. Interestingly, the phase of the circadian cycle seems to have a non-negligible effect on task engagement and ensuing performance (see Figure 1b). More precisely, mice tend to attempt less trials, abort trials more often and make more errors during daytime hours (between 8am and 6pm). Nevertheless, circadian effects are, by design, orthogonal with task factors. We will thus ignore circadian effects in the remainder of this paper. Because there is no clear way of interpreting aborted, unrewarded or unanswered trials, we consider these as invalid and exclude them from our analyses (about 22±7% of non-corrective trials).

Now, mice behavior in valid trials is itself the outcome of covert competing cognitive processes. In line with previous studies conducted in similar behavioral paradigms *(Ashwood et al., 2022; Roy et al., 2021)*, we classified each valid trial according to three candidate overt response patterns. First, a response was classified as “perseverative” if it was directed to the same side as the previous response. Second, a response was classified as “win-stay/lose-switch” (WSLS) if it repeated the previous response following a rewarded trial or switched sides following an unrewarded trial. Third, a response was classified as “cue-following” if it was directed toward the cued side. Importantly, these three criteria are partially orthogonal: a given response may satisfy any combination of them, including none. Figure 1c shows the evolving rates of these three response types averaged over mice, after having binned trials into bouts of 200 successive trials. During the training phase, mice progressively increased reward-seeking behavior (WSLS or cue-following responses) while reducing their reliance on the initially dominating perseverative behavior, eventually reaching the 66% accuracy criterion required to enter the main experimental phase. During the main experimental phase, both cue-following and WSLS response rates remained stable and significantly above chance level (one-sample *t*-test, p < 0.001). In principle, WSLS responses may originate from learned action-outcome associations. On the one hand, following an error, reinforcement learning mechanisms devalue the previously unrewarded action and thus facilitate a lose-switch response. Such prepotent, cue-independent, default response will, by design, achieve better-than-chance performance during corrective trials. On the other hand, at the onset of non-corrective trials (i.e. following success), the same learning mechanism will reinforce the previously rewarded action and thus promote a perseverative/win-stay response. However, since the reward location is now randomly sampled, independently of the mice’s previous actions, the prepotent default response will perform at chance level. Importantly, this does not imply that mice always override their default response during non-corrective trials. This can be seen in Figure 1d, which shows the evolving rates of both perseverative/win-stay and cue-following (correct) responses during non-corrective trials. Qualitatively, mice behavior during non-corrective trials (where default WSLS responses yield chance-level success) exhibits similar periods of improvement (pre-training phase) and plateau (main experimental phase) performance. But mice also maintain a stable and significant (one-sample *t*-test, p < 0.001) rate of default perseverative/win-stay responses. This suggests that, although mice mostly rely on cues to orient their decisions, they do not always override their default bias. Rather, as we will see, mice may regulate cognitive control on a trial-by-trial basis, eventually modulating conflict-induced interferences according to task demands.

### Computational model

Allocating control resources to instrumental information processing may bring reward but induces costs *(Kool et al., 2017; Otto et al., 2025; Shenhav et al., 2013a, 2017)*. So, what is the optimal resource allocation policy under possibly conflicting default and controlled responses? Our computational model relies upon three key assumptions: (i) prior to processing the visual cue, mice possess a prepotent default action bias, (ii) mice may override their default bias by engaging controlled processing of the visual cue, and (iii) mice regulate their control investment on a decision-by-decision basis, based upon an online cost/benefit arbitration.

Assumption (i) states that mice have attached unequal default values to candidate actions (here: touching either the left or right tactile screen). Such default values may derive from, e.g., reinforcement learning mechanisms, which would typically promote a perseverative/win-stay bias at the onset of non-corrective trials. Note that, formally speaking, such a default bias arises from a credit assignment error, whereby mice attribute rewards to candidate actions irrespective of whether they follow the cue or not. This error is a residue of pre-training, where mice progressively - though incompletely - discover that rewards can largely be attributed to cue-following actions. We refer the interested reader to Appendix 1 for computational modelling of the credit assignment problem and related analyses of pre-training data.

Assumption (ii) states that, as mice spend more time processing the visual cue, they increase the reliability of their perceptual evidence regarding the dominant orientation of the bars. This progressively renders the cue-following action more likely to bring reward, at a speed that depends upon decision difficulty. If the cue-dependent expected reward eventually reverses the default value difference, then mice override their default bias. We refer the interested reader to Appendix 2 for computational (Bayesian) modelling of the controlled attentional/perceptual processing of the visual cue.

Assumption (iii) requires mice to monitor and anticipate the net benefit of investing more cognitive control to sample the cue. On the one hand, the prospect of reward incentivizes mice to sample the cue and accumulate perceptual evidence. On the other hand, the growing cost of cognitive control will eventually overcompensate its expected benefit. One can show (see Appendix 3) that, at any point in time, there exists a critical evidence bound, above which the expected net benefit of investing further cognitive control will worsen. The optimal policy is thus to stop sampling the cue whenever the current perceptual evidence hits this bound. Because control cost progressively grows, critical evidence bounds decay over time. In addition, default action values alter the overall cost-benefit tradeoff of control. At the limit, the cost of control is incurred unnecessarily if cue processing ultimately confirms the default bias. This translates into asymmetrical evidence bounds for either confirming or overriding the default. More precisely, at any point in time, more perceptual evidence is required to stop and override the default action. As we will see, this eventually induces typical Stroop-like interference effects, i.e., response time and performance differences between trials where the cue is aligned with or opposed to the default response (“default-congruent” and “default-incongruent” trials, respectively). Importantly, the model also predicts the effect of motivational factors (namely: reward at stake and decision difficulty) on these interference effects. Note that the impact of default values effectively bridges previous motivational theories of cognitive control regulation *(Shenhav et al., 2013a, 2017)* and optimal accumulation-to-bound decision models *(Drugowitsch et al., 2012; Tajima et al., 2016)*. This represents the main computational contribution of this work, as it provides a unified framework from which we derive and test several novel predictions in the Results section.

## RESULTS

### Model predictions

The above decision control model predicts whether and how mice override their default response. In what follows, we disclose its underlying computational mechanisms.

During the early stages of cue processing, perceptual beliefs are very sensitive to random attentional sampling and thus unreliable (see Appendix 2), which may promote premature hits of the nearest (default) evidence bound. On average, quick responses are thus less likely to result in default overriding. In other words, the default bias is maximally expressed in rapid decisions. If the default response is not aligned with the visual cue (as is the case during *incongruent* trials), then this inflates the rate of (early) errors. This has two main consequences. First, incongruent trials typically yield lower performance than congruent trials (Fig. 2b). Second, decisions in incongruent trials are slower than in congruent trials, because, although most belief trajectories are attracted towards the correct bound in most cases, it takes more time on average to reach it (Fig. 2c). In brief, the model predicts that both performance and decision speed should worsen when the cue is not aligned with the default response.

**Fig. 2.**
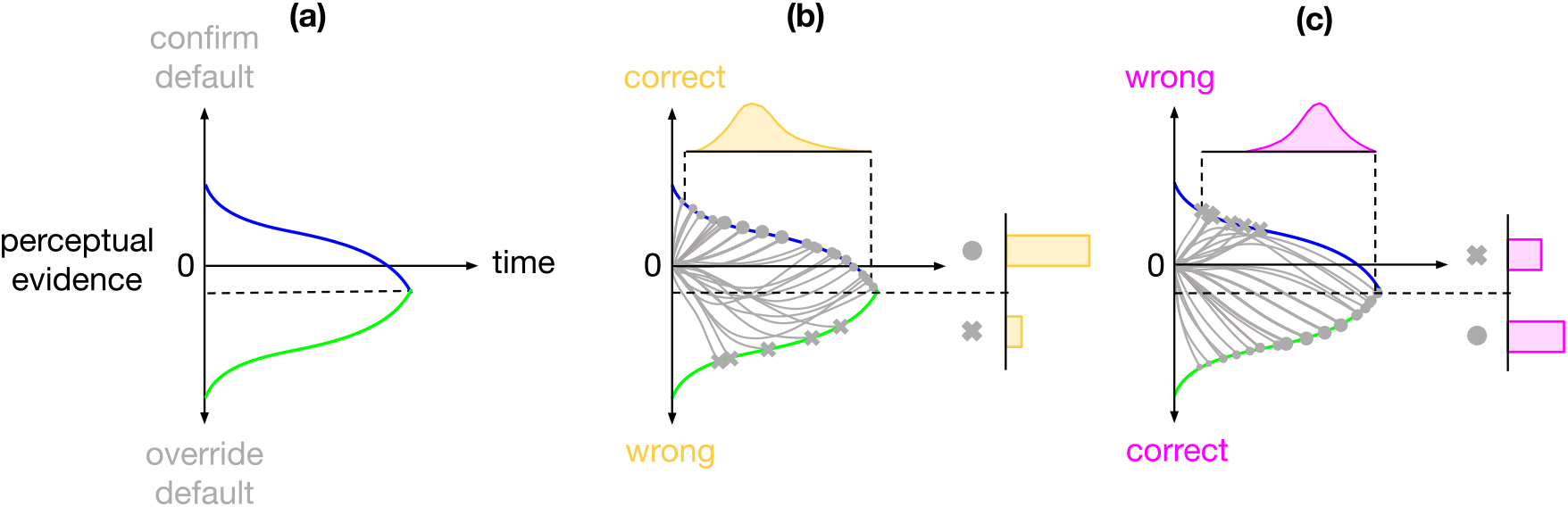
Default/automatic interference effects. **(a)** Dynamics of optimal evidence bounds plotted against decision time (blue: default-compliant, green: default-overriding). By convention, positive perceptual evidence confirms the default bias. In absolute terms, more evidence is required to trigger a default-overriding response (cf. black-dotted line: indecisive evidence level). **(b)** During default-congruent trials, the visual cue is aligned with the default bias. Therefore, default-following responses are correct, whereas default-overriding responses are wrong. In addition, most perceptual evidence sample paths eventually reach the (correct, more lenient) default-compliant bound, which induces a distribution of fast response times and mostly correct response outcomes (circle dots: correct responses, crosses: errors). This is because evidence trajectories that initially accumulate in favor of the incorrect choice generally have sufficient time to recover before crossing the incorrect bound. **(c)** Same as **a**, for default-incongruent trials. Here, the visual cue goes opposite to the default bias, hence default-following responses are wrong, whereas default-overriding responses are correct. Here, evidence sample paths take more time on average to reach the bounds (slower response times), though fewer of the initially erroneous trajectories have sufficient time to recover (lower rate of correct responses).

Now, reducing the reward at stake decreases the expected benefit of engaging control to process cue-related information. Similarly, increasing decision difficulty increases the amount of controlled processing required to reach any target evidence level. In both cases, the cost/benefit balance is less favorable, which eventually shrinks optimal stopping boundaries (see Appendix 3). This has three consequences. First, less evidence is integrated in the decision. As a result, controlled processing is less likely to override the default response on average. Hence, decreasing reward or increasing difficulty should decrease performance and increase the rate of default responses. Second, shrinking decision boundaries limits the opportunity for initially erroneous evidence trajectories to avoid premature default bound hits. Therefore, the performance loss in incongruent trials should increase as reward decreases or difficulty increases. Third, since premature boundary hits are more likely, the average difference in response times between congruent and incongruent conditions is reduced. Thus, in contrast to performance interference effects, response time interference effects should decrease when difficulty increases or reward decreases.

Finally, the model makes additional predictions regarding the relationship between response times on the one hand, and response rates (cue-following or default-following) or performance interference effects on the other hand. Recall that optimal evidence thresholds eventually collapse as time elapses. Thus, on average, later decisions are based on poorer evidence, i.e. the model predicts that the rate of correct (cue-following) responses decreases when response time increases. Importantly, the extra evidence level required to override the default bias also gradually decreases with decision time (see Fig. S4). Therefore, on average, the default bias becomes easier to override over time, i.e. the model predicts that the rate of default responses also decreases as response time increases. For the same reason, the performance advantage in congruent over incongruent trials is greatest for early decisions, when the default facilitation effect is strongest. In turn, the model also predicts that default interference effects in performance decrease as response time increases.

### Evidence for default/automatic interference effects

In what follows, we focus on non-corrective trials to preserve the orthogonality between default-following and cue-following responses. We also discard the hardest difficulty level (5:5) because the corresponding cue-following answer cannot be defined. For all statistical data analyses (here and below), we first estimated regression coefficients within each animal and then tested these coefficients against zero at the group level using one-sided t-tests (random effects analysis). Note that we do not apply correction for multiple comparisons across statistical tests here. This is because each test addresses a distinct, directional, *a priori*, model prediction, hence the ensuing family of tests is confirmatory rather than exploratory *(Armstrong, 2014; Hooper, 2025)*.

We begin by establishing behavioral evidence for conflict-induced interference effects. In non-corrective trials, mice’s default bias is to repeat the previous action (stay or win-stay response). We thus identify, for each non-corrective trial, the side of the default bias given the mice’s response at the previous trial. We then separated trials according to whether the visual cue was aligned (default-congruent trials) or not (default-incongruent trials) with the side of the default bias and quantified the corresponding rates of correct responses (Fig. 3a) and response time (Fig. 3b). Random effect analyses at the group level then allowed us to test the main effect of default congruency. First, we found that the rate of correct responses was significantly lower in incongruent than in congruent trials (p < 0.001). In addition, we found that response time was significantly higher in incongruent than in congruent trials (p = 0.032). In other words, both performance and decision speed worsen in incongruent trials as compared to congruent trials. Interestingly, performance and response time interference effects also shows a significant positive correlation across mice (r = 0.35, p = 0.04, one-sided t-test ; see Fig 3c). This is compatible with the notion that a common mechanism underlies both interference effects.

**Fig. 3.**
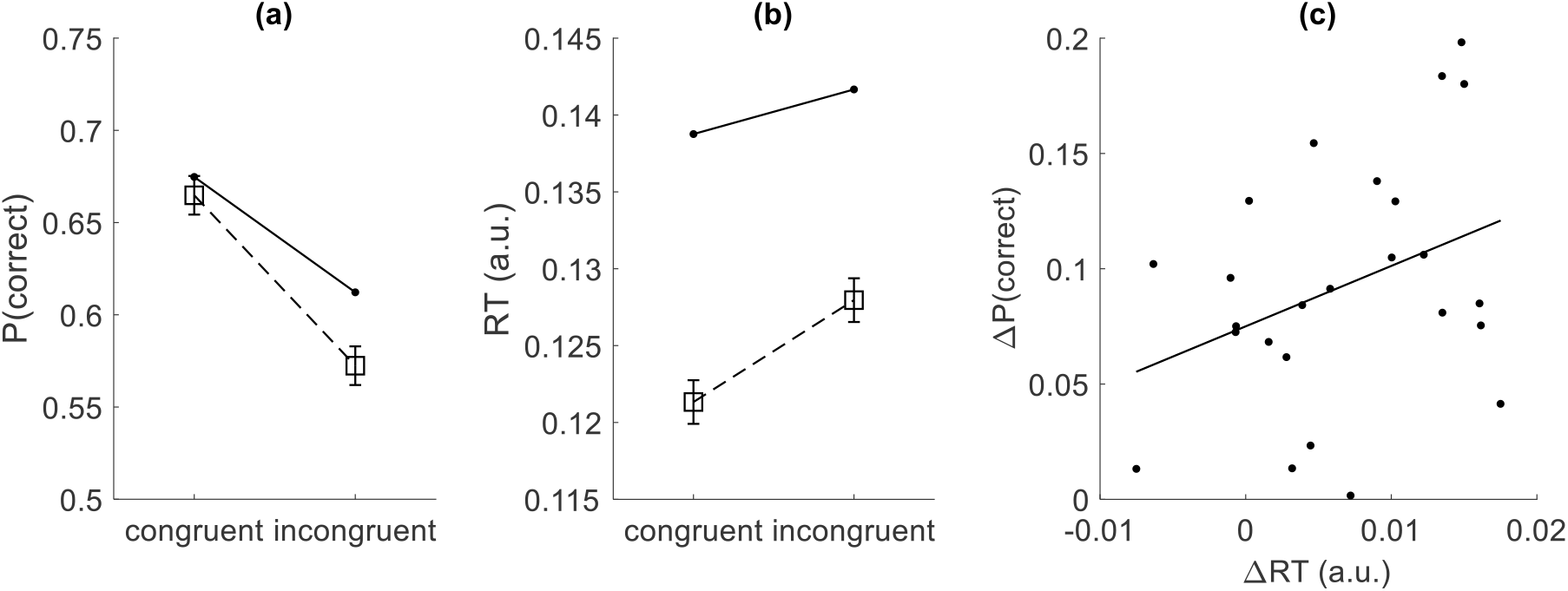
Default/automatic interference effects. **(a)** The average rate of correct responses is shown for both default-congruent and default-incongruent trials (plain lines: model predictions, dotted lines with squares: data). Error bars indicate 95% confidence intervals. **(b)** The average response time is shown for both default-congruent and default-incongruent trials (plain lines: model predictions, dotted lines with squares: data). **(c)** Performance interference (difference in correct response rates between congruent and incongruent trials) is plotted against response time interference (difference in response times), for each mouse (dots). The plain line shows the best linear fit.

### Motivational modulations of interference effects

We now ask whether and how the above interference effects are modulated by motivational factors that alter the cost–benefit balance of engaging controlled processing, in particular: reward at stake and decision difficulty. In brief, the model predicts that rendering the cost-benefit balance less favorable increases performance interferences and decreases response time interferences.

We tested the predicted modulation of performance interference effects by regressing binary performance data against trial congruence, difficulty (or reward), and their interaction for each mouse. Group-level analyses of regression coefficients revealed a significant positive interaction between difficulty and trial congruence (p = 0.006), as well as a significant negative interaction between reward and trial congruence (p = 0.037), in line with model predictions (see Fig 4a and 4b). To test the predicted modulation of response time interference effects, we regressed response times for each mouse against trial congruence, difficulty (or reward at stake), and their interaction. We found a significant negative interaction between difficulty and trial congruence (p < 0.001), as predicted by the model (Fig 4d). Note that although the trend is visible in the data (see Fig 4c), response time interference effects do not significantly increase when reward increases (p = 0.233).

**Fig. 4.**
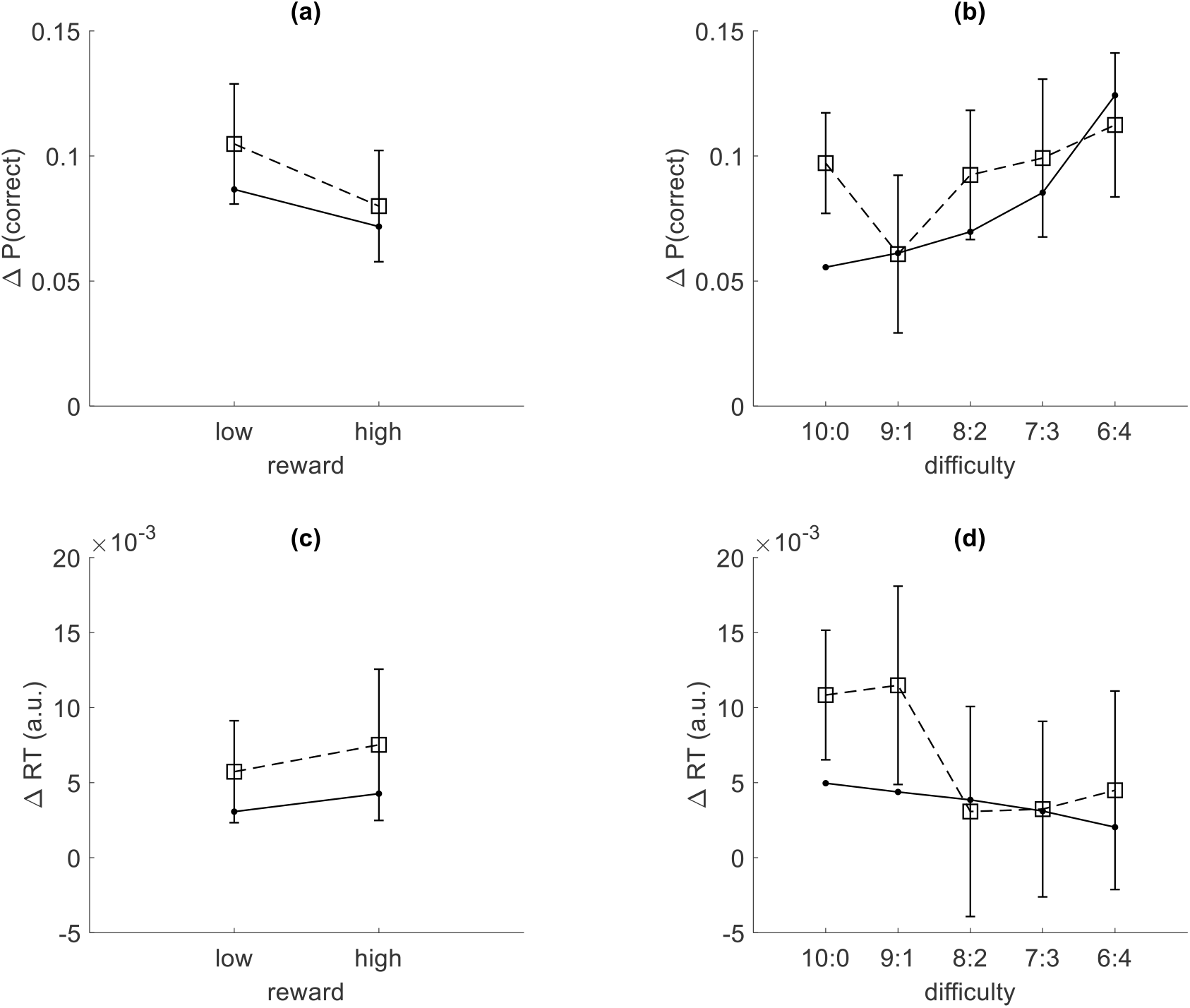
Motivational modulation of interference effects. **(a)** The average difference between correct response rates between default-congruent and default-incongruent trials is shown as a function reward (plain lines: model predictions, dotted lines with squares: data). Error bars indicate 95% confidence intervals. **(b)** Same as **a**, across difficulty conditions. **(c)** The average difference in response time between default-incongruent and default-congruent trials is shown as a function reward (plain lines: model predictions, dotted lines with squares: data). **(d)** Same as **c**, across difficulty conditions.

### Motivational modulation on the arbitration between response types

The model also predicts that motivational factors will shift the cost-benefit arbitration between default and cue-driven responses. Figure 5 below summarizes the ensuing analyses.

**Fig. 5.**
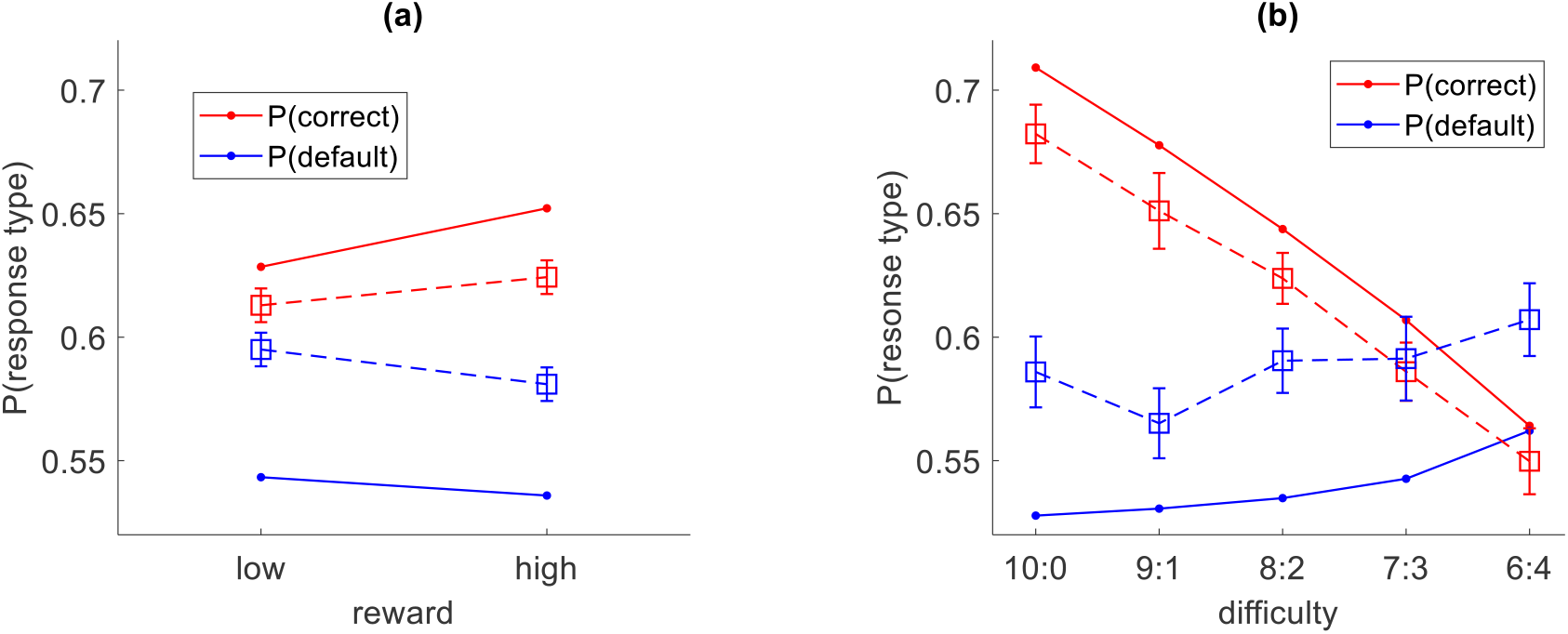
Motivational modulation of arbitration between response types. **(a)** The average rate of correct (red) and default (blue) responses are shown as a function reward (plain lines: model predictions, dotted lines with squares: data). Error bars indicate 95% confidence intervals. **(b)** Same as **a**, across difficulty conditions.

To test model predictions, we regressed the rate of cue-following and default-following responses onto difficulty and reward levels (and their interaction) within each mouse. As predicted by the model, group-level analyses of regression coefficients revealed that the rate of default-congruent responses significantly increases when reward decreases (p = 0.035, Fig 5a) and when difficulty increases (p < 0.001, Fig 5b). Also, as expected, performance significantly increases when reward increases (p = 0.04, Fig 5a) and when difficulty decreases (p < 0.001, Fig 5b).

### Relationship between decision time and interference effects

Finally, the model makes additional predictions regarding the relationship between response time on the one hand, and response rates (cue-following or default-following) or performance interference effects on the other hand. Figure 6 below summarizes the ensuing analyses.

**Fig. 6.**
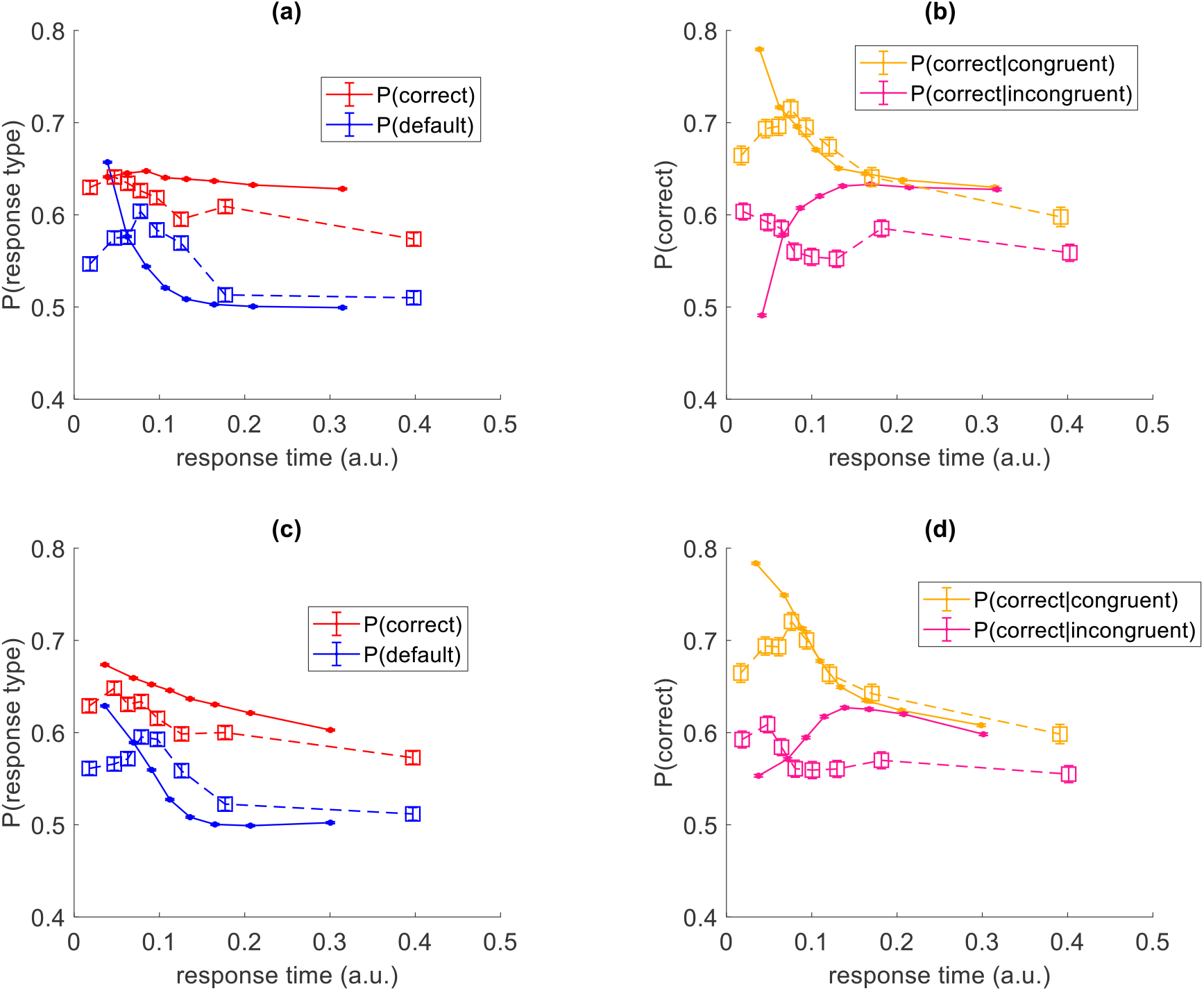
Linking response time to arbitration between response types and related performance interference effects. **(a)** The rate of both correct (red) and default (blue) responses (y-axis, plain lines: model predictions, dotted lines with squares: data) is potted against response time (x-axis, arbitrary units). Error bars indicate 95% confidence intervals. **(b)** The rate of correct responses (y-axis, plain lines: model predictions, dotted lines with squares: data) is potted against response time (x-axis, arbitrary units), for both default-congruent (orange) and default-incongruent (magenta) trials. Error bars indicate 95% confidence intervals. **(c**,**d)** Same as **a** and **b** (respectively), when controlling for reward and difficulty.

First, the model predicts that both performance and default response rates should decrease when response time increases. Thus, we regressed performance against response time for each mouse and tested regression coefficients at the group level. As expected, we observed that performance significantly decreases when response time increases (p = 0.002, Fig 6a). Importantly, this relationship remained significant after controlling for reward and difficulty (p < 0.001, Fig 6c). We then performed the same analysis for the rate of default responses. As predicted by the model, the rate of default responses significantly decreases when response time increases (p < 0.001, Fig 6a), and this relationship remained significant after controlling for reward and difficulty (p < 0.001, Fig 6c).

Second, the model predicts that performance interference effects decrease as response time increases. This prediction was tested by regressing performance against response time, trial congruence, and their interaction. Group-level analysis revealed a significant interaction between response time and trial congruence, indicating that interference effects on performance are indeed larger at shorter response times (p < 0.001, Fig 6b). Crucially, this interaction persisted after controlling for reward and difficulty (p < 0.001, Fig 6d).

## DISCUSSION

In this work, we have provided evidence that a single normative mechanism can account for the diverse behavioral signature of cognitive control regulation in mice. On the computational side of things, we have bridged existing motivational theories of cognitive control regulation *(Kool et al., 2017; Lieder et al., 2018; Shenhav et al., 2013a, 2017)* to optimal accumulation-to-bound decision models *(Bénon et al., 2024; Drugowitsch et al., 2012; Tajima et al., 2016)*. This enabled us to predict how motivational factors such as reward at stake and decision difficulty eventually modify the underlying cost/benefit tradeoff and thus impact the arbitration between default and controlled responses. On the behavioral side of things, we have validated all the ensuing model predictions, in the context of an experimental setup with high ecological validity. This opens an avenue for assessing the brain circuits that operate the arbitration between default and controlled decision processes.

To begin with, let us revisit our main computational contribution, which relates to the nature of default responses and the mechanisms by which they may override controlled processes. An important assumption is that, at the onset of a given decision, candidate actions have already been assigned distinct values. For example, these default values may originate from habit learning or other forms of reinforcement learning. This induces a prepotent default response - or rather, a default preference - whose key feature is that it is independent from the sort of contextual information that requires cognitive control to be processed (here: the visual cue). Importantly, default values alter the overall cost-benefit tradeoff of cognitive control, such that less evidence is required to trigger an action that confirms the default preference, all else being equal. This asymmetry in evidence bounds yields both a decision bias (i.e. ambiguous evidence tends to promote default-following responses) and a temporal facilitation effect (i.e., default-following responses tend to be quicker). Under the model, both effects are tied together and determined by the difference in default values, the importance of the decision, and the cost of control. This eventually explains established Stroop-like interference effects *(Cohen et al., 1990; Kalanthroff et al., 2018)*. Unspecific individual differences in these subjective features of the cost/benefit arbitration also explains why these two interference effects correlate across individuals *(Feltgen & Daunizeau, 2021; Lopez-Persem et al., 2016)*. Interestingly, the model also makes an entirely novel set of predictions, which relate to decision confidence reports. In brief, for any given response time, subjective reports of decision confidence should be lowest for default-following responses. Similarly, they should be higher in incongruent trials than in congruent trials. This confidence interference effect is not trivial, because it reverses the typical pattern of performance interference effects. We are currently testing these predictions in humans with a modified Stroop task.

Another important assumption of our work is that the arbitration between default and controlled processes is performed *in the frame of candidate actions*. What we mean here is that the decision system integrates default action values with cue-conditional expected rewards prior to triggering a choice. This maps onto modern neural net architectures of cognitive control, which rely on conflict monitoring within a layer of response units that integrate default-related and control-related processes *(Cohen et al., 1990; Musslick et al., 2016)*. Nevertheless, another alternative would be to perform the arbitration *in the frame of candidate processes*.

This implies deriving the optimal stopping policy by comparing the values of the preferred actions under default and controlled decision processes, respectively. Intuitively, this would gracefully implement the putative computational architecture of early opponent-process theories, whereby automatic (e.g. habitual or emotional) and controlled (goal-oriented or analytical) modes of behavioral guidance directly compete with each other *(Berridge, 2004; MacLeod, 1991)*. Although we do not report it here, we have derived the optimal policy under the latter scenario and analyzed its ensuing behavioral predictions. In brief, both computational scenarios tend to exhibit similar Stroop-like interference effects, except for response times. More precisely, arbitration in the frame of candidate processes yields no decision slowing when the default response is incongruent with the cue. We believe this disqualifies that particular – and otherwise reasonable – model variant.

Note that all model predictions shown in the Results section derive from a variational Bayesian parameter estimation approach, which we describe in Appendix 5. Importantly, adjusting the model parameters to match the sufficient statistics of the empirical data does not severely modify the qualitative predictions regarding the effects that we test and report in the main text. However, changing the model parameters that determine the cost-benefit tradeoff of cognitive control can impact the expected effect of some experimental factors. In particular, under the model, increasing decision difficulty can either tend to decrease (as we observe here) or to increase overall response time. The latter regime typically arises when the cost of control is low when compared to the reward at stake and/or when the system does not adapt its policy to difficulty. In such cases, response time variations across difficulty levels are mostly determined by the flattening of evidence trajectories, which will yield later bound hits on average (response slowing). This control regulation regime has been observed in humans, and is typically interpreted as a reactive and/or between-trial, control adjustment process *(Botvinick et al., 2001; Yeung & Cohen, 2006)*. This contrasts with motivational perspectives on cognitive control regulation, which rather highlight prospective, within-trial, cost-benefit optimization mechanisms *(Kool et al., 2017; Shenhav et al., 2013a, 2017; Westbrook et al., 2013)*. Having said this, for any given control cost, there exists a critical difficulty level above which the model would transition from a response-slowing to a response-speeding regime *(Bénon et al., 2024; Lee & Daunizeau, 2021)*. This also holds for response time interference effects. A possibility is that mice’s perceived difficulty was too high to observe the response-slowing regime of the model *(van Steenbergen et al., 2015)*. However, this has no bearing on the predicted variations in default-induced interference effects along response time variations, when controlling for difficulty (cf. Fig 6c and 6d).

Although our framework is fundamentally normative, it naturally accommodates empirical findings showing that psychobiological states such as stress, emotions, and fatigue alter decision makers’ tendency to rely on automatic/default responses. Within our framework, these effects need not invoke distinct decision mechanisms, but instead map onto changes in specific computational variables. For example, stress may induce an urgency signal *(Cisek et al., 2009)* that can be captured by shortening the temporal horizon. Similarly, fatigue effectively increases mental effort costs *(Pessiglione et al., 2025)* and emotions are known to bias reward-effort tradeoffs in accordance with their valence *(Heerema & Pessiglione, 2025)*. Importantly, this does not undermine the normative interpretation of the model, because variations in these internal states correspond to contextual alterations in the organism’s operating conditions. Consequently, the resulting shifts in the decision maker’s reliance on default biases may still be deemed adaptive from an evolutionary perspective *(de Kloet et al., 2005; Pessiglione et al., 2023; Verdeil, 2025)*.

This work has implications for the neural architecture underlying the control of goal-directed behavior. Previous empirical studies relying on accumulation-to-bound models provided evidence that the adjustment of decision thresholds may be operated by the basal ganglia, in response to inputs from the orbitofrontal cortex or OFC *(Frank, 2006)*. This is consistent with the fact that disruption of subthalamic nucleus or striatal activity may cause premature responding in humans *(Frank et al., 2007)* and promote default -e.g. perseverant-responses in mice *(Burguière et al., 2013)*. In contrast, exhaustive neuroimaging meta-analyses indicated that cost/benefit computations involved in cognitive control may engage the anterior cingulate cortex or ACC *(Shenhav et al., 2013b)*, which has known direct anatomical connections to the OFC and the basal ganglia *(Vogt & Paxinos, 2014)*. This raises the questions of whether and how ACC contributes to the control of decision making. Although the present model does not uniquely identify the neural implementation of these computations, it provides a computational framework for testing whether ACC-dependent control signals modulate decision thresholds and/or control costs. We are currently addressing this issue using chemogenetic manipulations of ACC activity in mice.

This work also has implications for neuropsychiatric conditions marked by aberrant allocation of cognitive control, most directly obsessive-compulsive disorder (OCD). OCD is characterized by intrusive, perseverative behaviors that persist despite conflicting goal-directed information. Importantly, converging evidence for the underlying pathophysiological mechanism already exists: disruption of orbitofrontal-striatal and subthalamic pathways promotes compulsive-like behavior in mouse models *(Burguière et al., 2013)*. Under our model, this would qualify as excessive reliance on premature default-biased responding, which may arise from reduced sensitivity to reward, heightened sensitivity to the cost of cognitive control, or miscalibrated beliefs about the expected benefit of cognitive control. Our task, itself adapted from a cross-species compulsivity assay *(Benzina et al., 2021)*, offers a direct route to testing whether OCD-relevant circuit manipulations shift the default/controlled arbitration boundary predicted by the model. Moreover, the same framework extends to other conditions marked by dysexecutive symptoms, such as treatment-resistant depression (where loss of cognitive, affective, and behavioral control frequently cooccur) and ADHD (where impulsive/premature responding is a hallmark symptom). Thus, the computational approach presented here may help decompose overtly similar clinical phenotypes into distinct, mechanistically separable computational profiles.

**APPENDICES**

## Appendix 1 The credit-assignment problem of pre-training

To complete the last pre-training phase of the experiment, mice need to reach a 66% correct response rate, where the correct response is given by the cue. At this point, they need to have partly solved the underlying credit assignment problem. From the mice’s perspective, that latter can be stated as the following question: are rewards contingent on action side (default attribution), or are they contingent on the congruence of cues and action (hereafter: cue-related attribution)? Formally speaking, this can be reframed as a Bayesian inference problem. Let *m* ∈ {0,1} be a binary variable that flags whether the default (*m* = 0) or the cue-related attribution (*m* = 1) is true. In what follows, we refer to it as the *credit variable*. Let *a*_*t*_ ∈ {0,1}, *c*_*t*_ ∈ {0,1} and *r*_*t*_ ∈ {0,1} be the side of the chosen action (0: left, 1: right), the side indicated by the visual cue (0: left, 1: right) and the obtained reward (0: no reward, 1: reward) at trial *t*, respectively.

Under the default attribution (*m* = 0), the likelihood of obtaining the reward can be written as the following Bernoulli distribution:

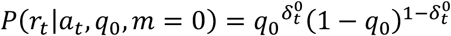

where *q*_0_ = *P*(*r*_*t*_ = 1|*a*_*t*_ = 1, *m* = 0) is the probability of obtaining the reward when choosing *a*_*t*_ = 1 (e.g., picking the left side) and 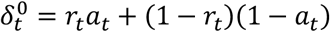 is a binary variable that flags if the obtained reward is congruent with the action side 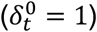 or not 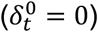.

Under the cue-related attribution (*m* = 1), the reward likelihood is rather given by:

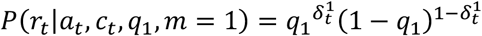

where *q*_1_ = *P*(*r*_*t*_ = 1|*a*_*t*_ = *c*_*t*_, *m* = 1) is the probability of obtaining the reward when choosing *a*_*t*_ = *c*_*t*_ (i.e. following the cue), and 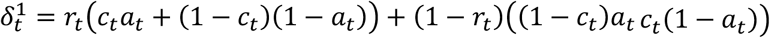 is a binary variable that flags if the obtained reward is congruent with following the cue 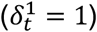 or not 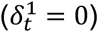. Note that, in our experimental setting, 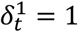 by design (i.e. the event 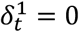 is impossible). This simplifies the above reward likelihood as follows: *P*(*r*_*t*_|*a*_*t*_, *c*_*t*_, *q*_1_, *m* = 1) = *P*(*r*_*t*_|*q*_1_, *m* = 1) = *q*_1_.

Solving the ensuing credit assignment problem reduces to updating one’s posterior belief *P*(*m, q*_0_, *q*_1_|*r*_1:*t*_, *a*_1:*t*_, *c*_1:*t*_) about the unknown credit variable *m* and related probabilities *q*_0/1_ given the sequence of past actions, cues and rewards (up to trial *t*):

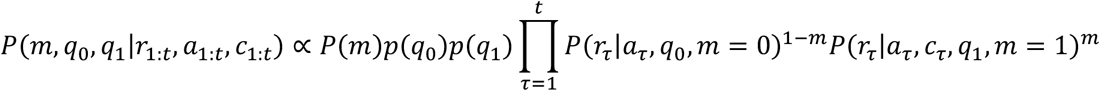

where *P*(*m*), *p*(*q*_0_) and *p*(*q*_1_) are prior beliefs about the unknown variables of the credit assignment problem (at the onset of pre-training).

A variational Bayesian approach (Blei et al., 2017; Daunizeau, 2018; Friston et al., 2007) to this problem yields the following approximate marginal posterior distributions for the credit variable *m* and related probabilities *q*_0/1_:

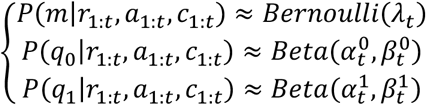

whose sufficient statistics are iteratively updated as follows:

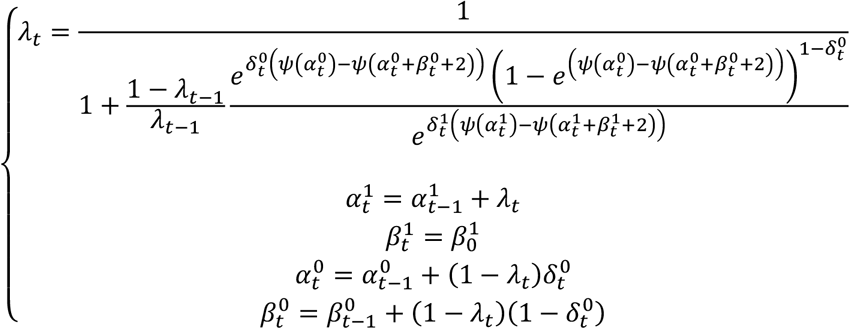

where *ψ*(·) is the digamma function and we have used the fact that 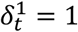 at all trials.

Here, *λ*_*t*_ = *P*(*m* = 1|*r*_1:*t*_, *a*_1:*t*_, *c*_1:*t*_) is the posterior belief on the credit variable at trial *t*, i.e. the subjective probability estimate that rewards are contingent upon cue-following actions (given past rewards, actions and cues up to trial *t*). Note that, at trial *t*, the agent’s posterior estimate of probabilities *q*_0_ and *q*_1_ are given by 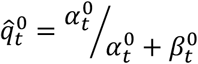 and 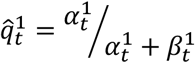, respectively.

At trial *t* + 1, the agent’s tendency to choose either *a*_*t*+1_ = 1 or *a*_*t*+1_ = 0 relies upon her conditional reward expectation. Now, the probability of obtaining the reward depends upon the cue, the agent’s posterior belief, and the chosen action side:

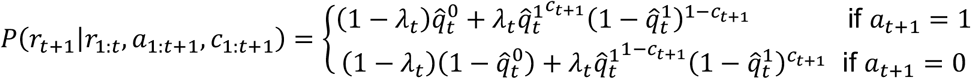

The agent’s behavior is driven by two competing forces, which directly derive from either the default or the cue-following attribution. The balance between these two forces is progressively modified as the agent updates her posterior belief *λ*_*t*_ on the credit variable. More precisely, as the credit assignment converges onto the cue-related attribution (i.e. as *λ*_*t*_ → 1), the agent’s behavior is increasingly determined by the cue.

Let Δ*Q*_*default*_ and Δ*Q*_*cue*_ be the reward expectation difference between actions (here: a = 1 and a = 0) under each candidate attribution hypothesis (i.e. *m* = 0 or *m* = 1):

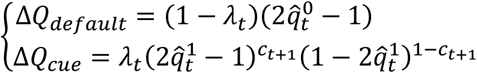

To compare the above Bayesian credit-assignment model to mice’s choices during pre-training, we assume that the probability that mice choose action *a*_*t*+1_ = 1 is given by:

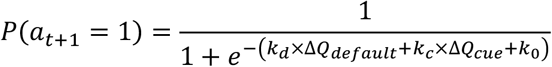

where *k*_*d*_ and *k*_*c*_ weigh the default and cue-related behavioral forces, and *k*_0_ is some additional side bias. We rely on the VBA toolbox *(Daunizeau et al., 2009, 2014)* to adjust the unknown parameters *k*_0_, *k*_*d*_ and *k*_*c*_ (as well parameters 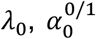 and 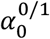 that quantify mice’s prior beliefs) to observed action side sequences during pre-training, separately for each mouse. The results of this analysis are given in Figure S1 below.

**Fig. S1.**
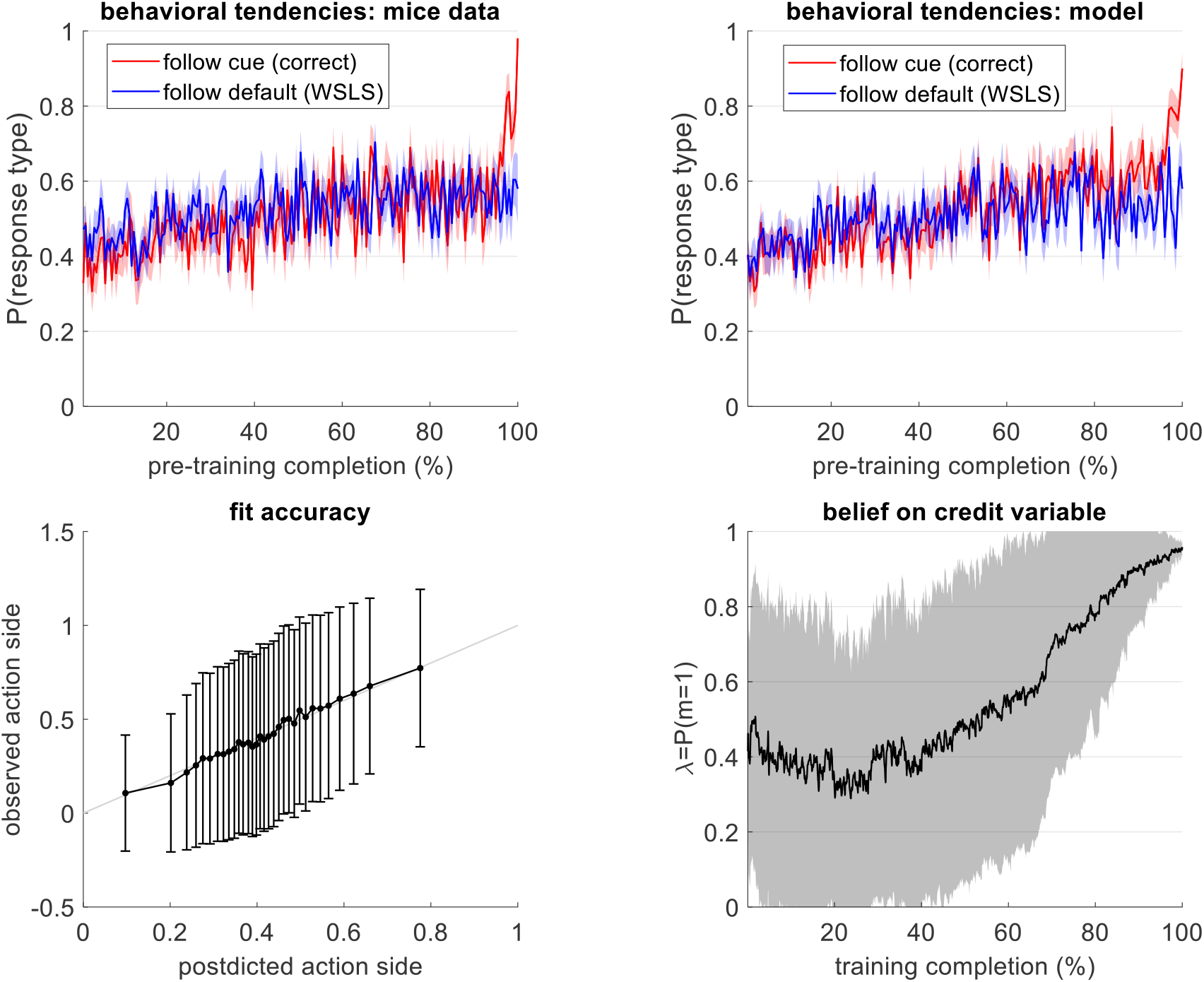
Results of pre-training data analyses under the Bayesian credit-assignment model. Upper-left panel: the rate of mice choices (y-axis) following either the cue (red) or the default (blue) is plotted against pre-training completion (x-axis: in % of pre-training trials until the switch to the main experimental phase). Upper-right panel: same, as postdicted by the Bayesian credit-assignment model. Lower-left panel: the observed rate of mice’s action side (y-axis) is plotted against the postdicted action side (x-axis). Lower-right panel: the dynamics of the posterior belief *λ*_*t*_ on the credit variable is plotted against pre-training completion. In all panels, shaded areas and error bars indicate the standard deviation across mice.

To begin with, we asked whether and how mice modify their behavioral responses as pre-training unfolds. To do this, we flagged each trial according to whether the corresponding action side was congruent with the visual cue or with a default (win-stay/lose-switch) heuristics. We then interpolated mice-specific sequences of pre-training trials onto a common “training completion” period that starts and ends at the onset and offset of their respective pretraining session. The average of the ensuing cue-following and default-following behavioral dynamics can be eyeballed on the upper panels of Figure S1. As pre-training unfolds, mice progressively increase their tendency to follow the cue, until this eventually dominates over their default behavioral tendency. The Bayesian credit-assignment model reproduces this pattern, although it seems to slightly over-anticipate the dominance of cue-following responses.

The lower-left panel of Figure S1 depicts the trial-by-trial fit accuracy of the Bayesian credit-assignment model. In quantitative terms, the model explains 4.9% ±0.5% of action side trial-by-trial variance. Note that R^2^ in the low single digits is unsurprising because irreducible trial-level noise caps achievable R^2^ well below 100% for single-trial binary choice data. This simply means that the model ignores other behavioral determinants that may induce fluctuations in mice’s choices. These include, but are not limited to, perseverative behavior, confusion about the visual cue, forgetting effects, satiety, etc. Nevertheless, the model is clearly able to predict mice’s choices better than chance. This is remarkable, given the huge number of pre-training trials (n=1327 ±94) when compared to the simplicity of the model (only 8 parameters).

Finally, we looked at the average dynamics of the estimated posterior belief *λ*_*t*_ on the credit variable, under the credit-assignment model. In brief, mice seem to remain uncertain in their reward attribution (*λ*_*t*_ ≈ 0.5) until about 50% of pre-training completion. They then tend to favor the cue-related attribution, until they reach *λ*_*t*_ ≈ 0.95 at the time when cue-following responses start dominating their behavior. Notably, under the credit-assignment model, mice did not completely eliminate the default attribution by the end of the pre-training phase (i.e. at the onset of the main test phase). This explains why mice exhibit a default bias during the main experimental phase.

## Appendix 2 Controlled processing of the cue

In our experimental setting, visual cues correspond to a set of 10 lines that are either vertical or horizontal (e.g. a 3:7 horizontal to vertical lines partition). The correct response depends on whether there are more horizontal than vertical lines or not, and we directly manipulate the difference between the number of vertical and horizontal lines (hereafter: Δ*n*).

To properly orient their response in accordance with the cue, mice need to infer its dominant orientation (vertical versus horizontal), under the constraint that her visual field of view spans a restricted portion of the cue. Mice thus sample the visual cue in a sequential manner, by actively shifting their attentional focus. Each foveation effectively extracts a spatially restricted field of view that is processed by the visual system. For the sake of simplicity, we assume that the visual processing of the attended field of view can be described in terms of a simple Fourier decomposition along both vertical and horizontal frequencies *(Niell & Stryker, 2008)*.

Let *y*_*t*_, *B*_*h*_ and *B*_*v*_ be the vectorized restricted field of view at decision time *t* and standard Fourier basis function sets over horizontal and vertical frequencies, respectively. At each time point within a decision trial, the visual system extracts the energy difference Δ*e*_*t*_ between horizontal and vertical frequencies: Δ*e*_*t*_ = *y*_*t*_^*T*^(*B*_*h*_*B*_*h*_^*T*^ − *B*_*v*_*B*_*v*_^*T*^)*y*_*t*_. The ensuing sequence of sampled differential energies Δ*e*_*t*_ evolves in a pseudo-random manner, as the attentional trajectory samples distinct portions of the visual cue. The first two moments of the induced attentional sampling distribution of differential energies Δ*e*_*t*_ are exemplified in Figure S2 below.

As expected, the sample mean *E*[Δe] is directly proportional to Δ*n*. In addition, the standard deviation *std*[Δe] increases when |Δ*n*| → 0. Taken together, this means that the signal-to-noise ratio *SNR* = |*E*[Δe]|/*std*[Δe] of the attentional processing of the visual cue increases when |Δ*n*| increases, or equivalently, when decision difficulty decreases (see Fig. S2d). Of note, the sample mean *E*[ē] of the total energy *ē*_*t*_ = *y*_*t*_^*T*^(*B*_*h*_*B*_*h*_^*T*^ + *B*_*v*_*B*_*v*_^*T*^)*y*_*t*_, i.e. stimulus luminance, also follows decision difficulty (see Fig. S2c). Since the total energy *ē* has a much higher SNR than the differential energy Δe, it may signal decision difficulty much before the correct response can be identified (at the limit, at cue onset).

**Fig. S2.**
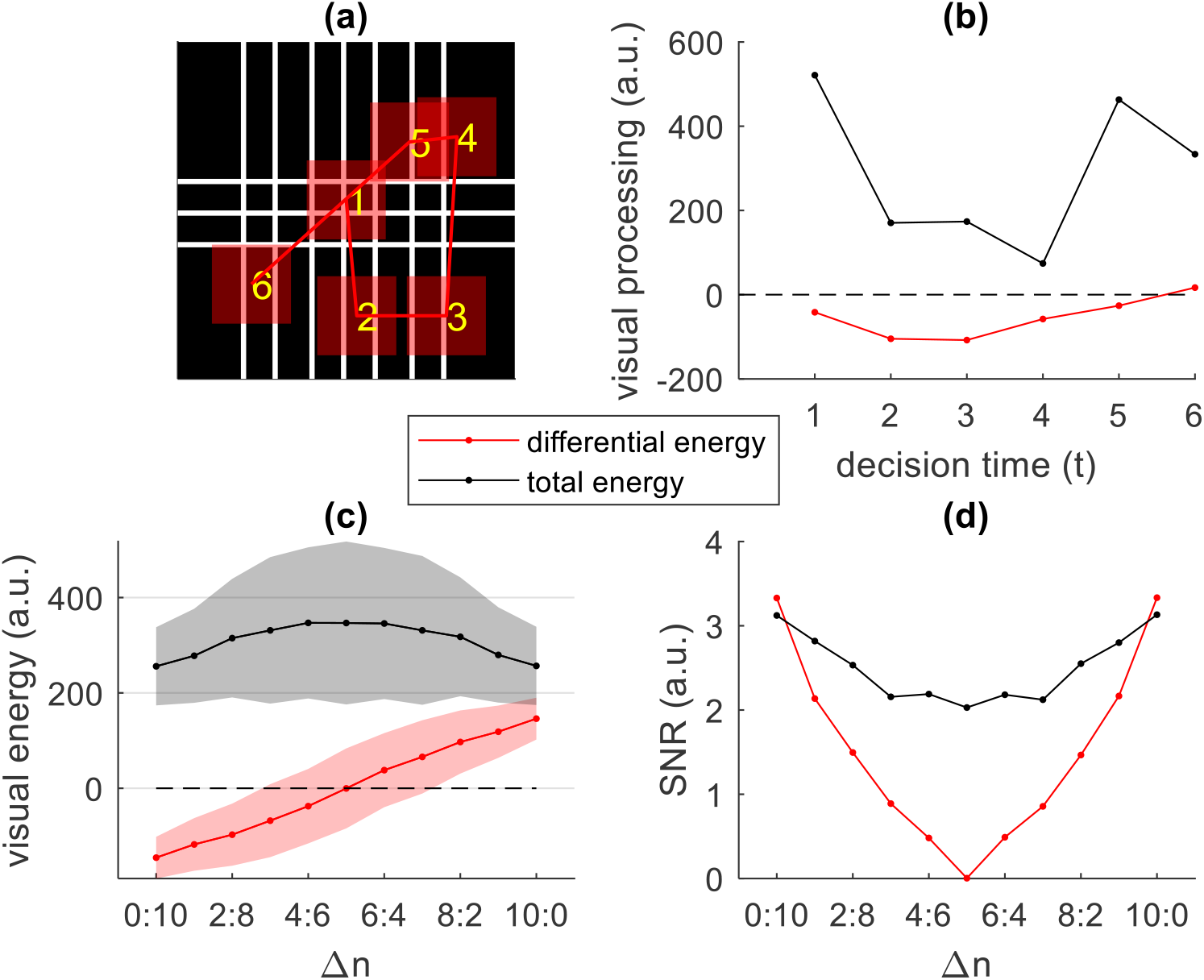
Attentional sampling model of the visual cue. **(a)** An example visual cue of moderate difficulty (3:7) is shown, along with a sequence of six random attentional foveations that span restricted portions of the cue (red shaded squares). **(b)** The ensuing sequence of sampled differential (Δ*e*_*t*_, red line) and total (*ē*_*t*_, black line) energies that are extracted by the visual system are plotted as a function of decision time. **(c)** The sample mean (10,000 Monte-Carlo samples) of both differential energy (*E*[Δe], red) and total energy (*E*[ē], black) are plotted as a function of Δ*n*. Shaded areas indicate the corresponding standard deviations *std*[Δe] and *std*[ē]. **(d)** The signal-to-noise ratio of differential energy (red line) and total energy (black line) are plotted as a function of Δ*n*.

At this point, we need to describe how the noisy trajectory of differential energy Δ*e*_*t*_ extracted by the visual system is then processed by the decision system to extract decision-relevant information. Formally, this can be framed as a Bayesian denoising process that aims at estimating the dominant orientation of the bars. Without loss of generality, we assume that, at the onset of a decision trial, the decision system starts with a non-informative prior belief about the visual cue:

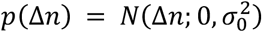

where 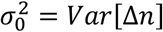 is the prior magnitude of Δ*n* and *E*[Δ*n*] = 0 a priori. We also assume that the decision system receives, at each time *t* after cue onset, a noisy, Gaussian-distributed (under the central limit theorem over the many contributing frequency components), evidence signal *S*_*t*_ ∝ Δ*e*_*t*_ centered on Δ*n* (i.e. *E*[S] = Δ*n*, cf. Fig. S2c):

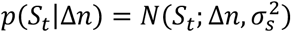

where 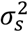 is the noise variance and the pseudo-random noise (*S*_*t*_ − Δ*n*) is induced by the arbitrary attentional trajectory spanning the visual cue (cf. Fig. S2b). Bayes’ rule then yields the posterior belief about Δ*n* after having observed the sequence of noisy evidence *S*_1:*t*_ up to time *t*:

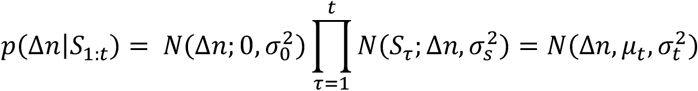

where the mean *μ*_*t*_ and variance 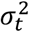 of the posterior belief *p*(Δ*n*|*S*_1:*t*_) at time *t* are given by:

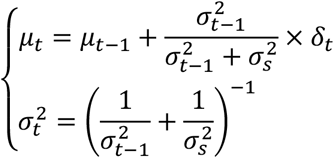

Here, *δ*_*t*_ = *S*_*t*_ − *μ*_*t*−1_ is the prediction error of the Bayesian agent.

This prediction error behaves as a stochastic variable driven by the arbitrary attentional sampling of the visual cue. This implies that the agent’s posterior mean *μ*_*t*_ will follow a Markovian random walk. Moreover, the Bayesian agent can predict, with some uncertainty, how her own belief will evolve over time. In fact, under her prior assumptions, she can form a posterior predictive belief about *δ*_*t*_, with first and second order moments given by:

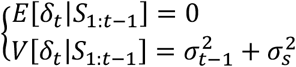

In turn, this defines a predictive transition density 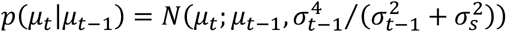 at time *t* − 1. That *E*[*μ*_*t*_|*μ*_*t*−1_] = *μ*_*t*−1_ means that the agent does not know in which direction her belief will change (this is because *E*[*δ*_*t*_|*S*_1:*t*−1_] = 0). However, the expected magnitude of this change is known and given by 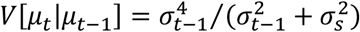.

Now, at time *t*, the Bayesian agent can guess the reward location from her uncertain belief 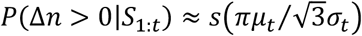 about the dominant orientation of the bars, which is the only decision-relevant information:

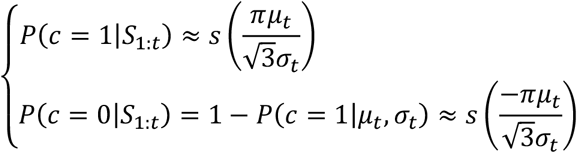

where *s*(*x*) = 1/(1 + *e*^−*x*^) is the standard sigmoid mapping, *c* = 1 (resp. *c* = 0) means that the cue indicates the left (resp., the right) side, and we have assumed that the agent has converged on the solution to her credit-assignment problem during the preceding pre-training stage.

Note that, formally, the sign of *μ*_*t*_ is a matter of convention, since the relationship between the dominant orientation of the bars and the rewarded side is randomized across mice. Since the direction of cues are randomized, we assume that the Bayesian agent starts with a non-informative prior (*μ*_0_ = 0). Should a decision be based uniquely upon this prior belief, its success rate would be at chance level (50%). Now, the sampling noise variance 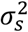 increases with trial difficulty, which can be rapidly proxied by the average stimulus luminance (*ē*_*t*_). This suggests that mice may prospectively respond to decision difficulty by adapting their expected noise variance 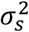 at the onset of each decision trial. We exploit this property in the decision model below.

## Appendix 3 Optimal stopping of the controlled cue processing

At this point, we ask when the agent should stop processing the cue and commit to a decision, given the expected costs and benefits of cognitive control. This optimal stopping problem is a specific instance of a Markov decision process *(Papadimitriou & Tsitsiklis, 1987)* that can be solved using dynamic programming.

Recall that the agents aims to maximize her expected net benefit *E*[*V*_*t*_ − *C*_*t*_], where *V*_*t*_ is the value of deciding at time *t* and *C*_*t*_ is the cost of controlled processing of the cue until time *t*. Under the assumption that the solution to the credit-assignment problem has converged, the value *V*_*t*_ of deciding at time *t* is:

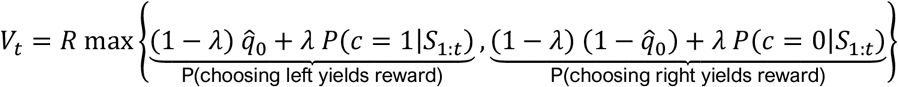

where *λ* is the agent’s belief that rewards are contingent upon the congruence of cues and actions (cf. Appendix 1), 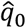 is her belief that choosing the left side will yield reward under the default attribution (for a given trial, 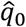 is constant), and *R* is the reward at stake.

At this point, we note that it does not really matter how *λ* and 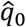 are set, as long as they induce a default action bias. By convention, we define the sign of cue-related evidence in relation to the side that is favored under the default bias, i.e. *μ*_*t*_ > 0 if the perceived cue is aligned with the default and *μ*_*t*_ < 0 otherwise. The value *V*_*t*_ can thus be rewritten as:

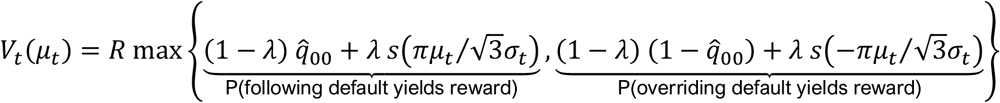

where 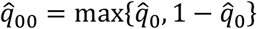 is the agent’s belief that she will get the reward under the default attribution, and we have rendered the dependence in *μ*_*t*_ explicit. Note this is a different sign convention than in Appendix 2, chosen here to align with the default bias for clarity of exposition.

Without loss of generality, we assume that *C*_*t*_ is an increasing function of controlled processing time *t (Otto et al., 2025)*, e.g. 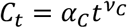, where *α*_*C*_ and *ν*_*C*_ determine the cost dynamics. In turn, even in open-ended decision problems, there exists a maximum processing time *H*, beyond which *V*_*t*_(*μ*_*t*_) − *C*_*t*_ is smaller than the net value of investing no control *V*_0_(0), no matter what. This happens at the latest when 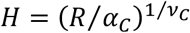. Strictly speaking, this is an upper bound on the maximum processing time. Nevertheless, we refer to it as the *temporal horizon* of the stopping problem, because overriding default biases is expected to bring no benefit beyond *H*. We note that, when adjusting model parameters to empirical data (see Appendix 5), the temporal horizon happen to lie beyond the response time limit that is imposed to mice in the task.

In what follows, we consider that the horizon is divided into *T* discrete time intervals, indexed by *t*. Let *a*_*t*_ ∈ {0,1} be the binary decision to continue (1) or stop (0) at processing time *t*. We are looking for the optimal stopping policy 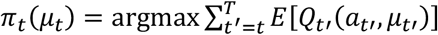, where *Q*_*t*_(*a*_*t*_, *μ*_*t*_) is the net benefit of controlled processing at time *t*:

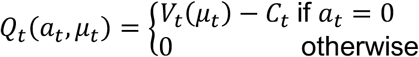

and the optimization is performed w.r.t. the sequence of all future actions {*a*_*t*_, *a*_*t*+1_, …, *a*_*T*_}. That the above net benefit is not realized until the agent stops and commits to a choice is a mathematical convenience: it just defers the payoff to whichever future timestep the agent stops at, so the backward induction scheme below correctly accumulates exactly one payout per trial.

By definition, the optimal stopping policy is to stop (*π*_*T*_(*μ*_*T*_) = 0) when the agent reaches the trial’s time horizon *T*. The ensuing net benefit is *Q*_*T*_(0, *μ*_*T*_) = *V*_*T*_(*μ*_*T*_) − *C*_*T*_, and which side the agent picks depends upon the evidence level *μ*_*T*_.

At time *T* − 1, the optimal policy is *π*_*T*−1_(*μ*_*T*−1_) = 0 (stop) if *Q*_*T*−1_(0, *μ*_*T*−1_) > *E*[*Q*_*T*_(0, *μ*_*T*_)|*μ*_*T*−1_], and *π*_*T*−1_(*μ*_*T*−1_) = 1 (continue) otherwise, where the expectation is taken under the transition density *p*(*μ*_*T*_|*μ*_*T*−1_). This transition density is derived in the Appendix 2 above. Importantly, there are two stop/continue frontiers (see Figure S3), located at two critical evidence bounds 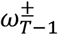, such that 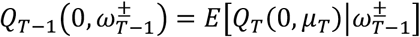.

Now, the *optimal net benefit* 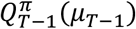, i.e. the net benefit obtained under the optimal stopping policy, is thus given by: 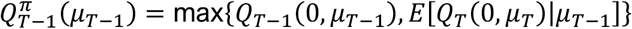. At time *T* − 2, the optimal policy is *π*_*T*−2_(*μ*_*T*−2_) = 0 if 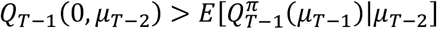 and *π*_*T*−2_(*μ*_*T*−2_) = 1 otherwise. The ensuing optimal net benefit is 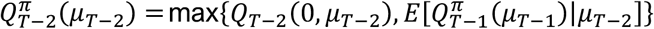.

This recursive induction scheme repeats backwards in time. Let 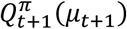 be the optimal net benefit at time *t* + 1. At time *t*, the optimal stopping policy is given by:

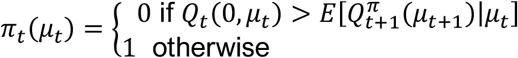

and the ensuing optimal net benefit is 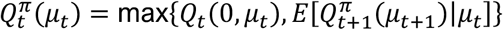, As before, two critical evidence bounds 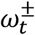 separate stop/continue decisions. This means that the optimal policy can be rewritten as:

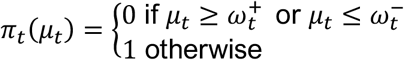

and which threshold is hit determines whether the agent overrides 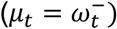 or complies with 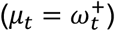 the default. If 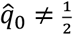, then 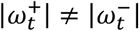, i.e. the evidence magnitude that is required to trigger a decision depends upon whether the signed evidence is aligned with the default bias. More precisely, 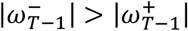, i.e. overriding the default bias demands more evidence, and the extra evidence required increases with 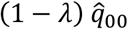 and decays over decision time (see Figure S3d below). The asymmetry between default-compliant and default-overriding evidence bounds is the reason why the model produces interference effects in response time and performance.

This concludes the derivation of the optimal policy.

In summary, a system trying to best balance the expected benefit and costs of controlled information processing will converge on a stopping policy that becomes more lenient as within-decision time elapses. Moreover, the optimal stopping policy demands more evidence to override a default response, and the extra evidence required to trigger a decision decreases over time.

**Fig. S3.**
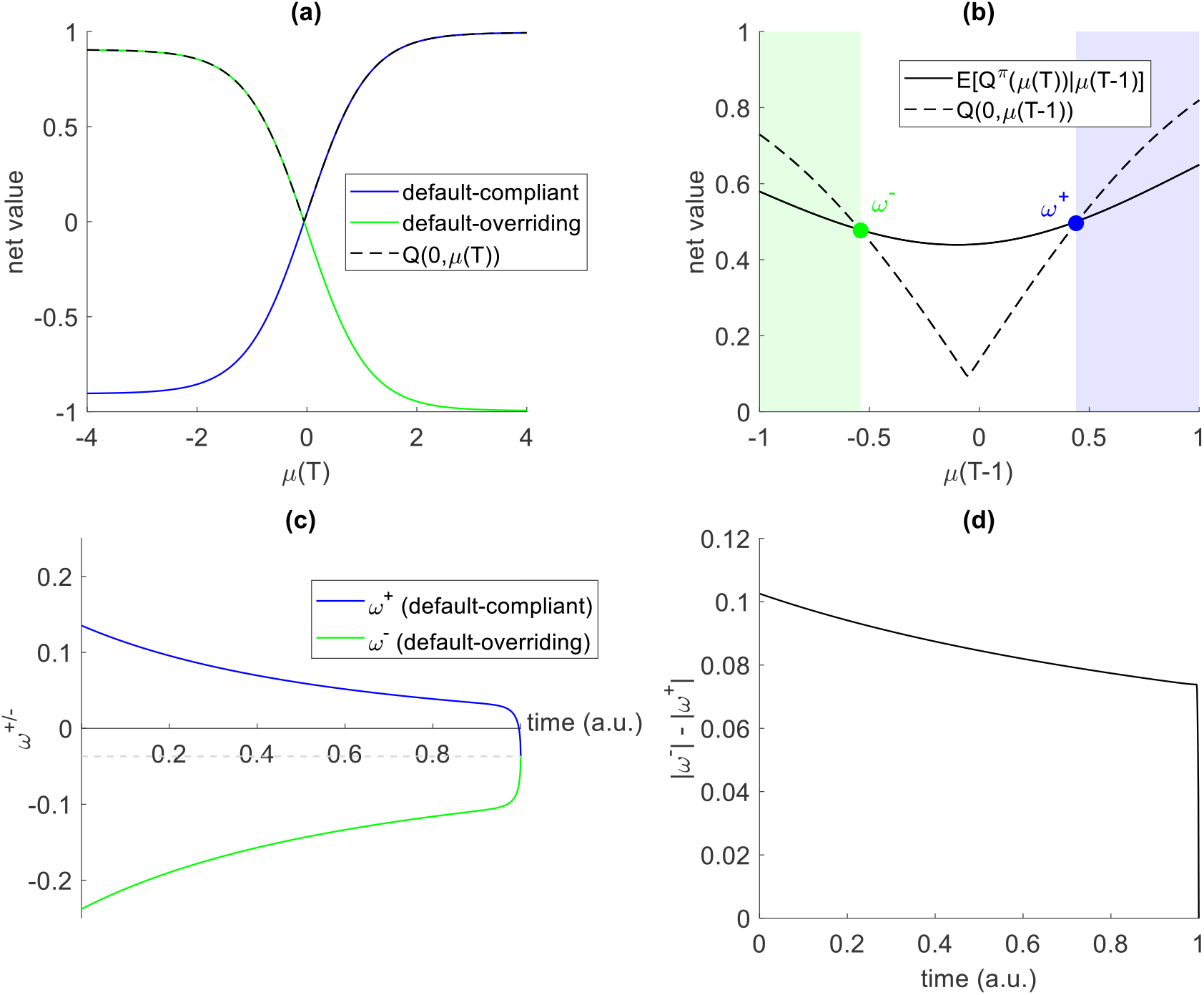
Optimal stopping policy. **(a)** At the temporal horizon, the net value of complying with the default (blue), overriding the default (green), and ensuing net benefit *Q*_*T*_(0, *μ*_*T*_) (dotted black) are plotted as a function of *μ*_*T*_. **(b)** One time step before the temporal horizon, the net value of stopping now *Q*_*T*−1_(0, *μ*_*T*−1_) (dotted black line) and the expected optimal net benefit *E*[*Q*_*T*_(0, *μ*_*T*_)|*μ*_*T*−1_] (plain black line) are plotted as a function of *μ*_*T*−1_. Crossing points show the two critical evidence bounds (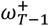 and 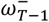), which would trigger a decision to stop. **(c)** The dynamics of critical evidence bounds 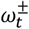 are plotted as a function of decision time *t* (normalized by the temporal horizon). The dotted black line indicates the indecisive evidence level, i.e. the evidence level at which both optimal evidence bounds eventually converge to. **(d)** The difference 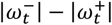 in the evidence magnitude required to trigger a decision is plotted as a function of decision time *t*.

Of course, the optimal policy, and hence the optimal evidence bounds, depend upon the model parameters (in Fig S3c, all parameters were set to 1 except for *λ* and 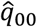, which were set to 0.95). For example, raising the reward at stake or decreasing the cost of control will expand both evidence bounds *(Bénon et al., 2024)*. In addition, decision difficulty has an impact through the moments of the perceptual signal (in particular: the noise variance *σ*_*s*_ ). This is because increasing the noise variance dampens the expected benefit of control, which eventually shrinks evidence bounds. Finally, the strength 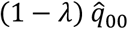 of the default bias will determine the asymmetry between default-compliant and default-overriding bounds.

Importantly, all model predictions reported in the main text were jointly derived using a single parameterization of the model (see Appendix 5 below).

## Appendix 4 Additional analyses of responses times

Orthogonally to the Stroop-like interference effects on response time (cf. Fig. 3), the model predicts that, on average, decisions that result in an error are slower than correct decisions. This is because late decisions are based on perceptual evidence of lower quality (cf. decaying optimal evidence bounds). Similarly, the model predicts longer response times for responses that do *not* follow the default response than for those that do. This follows from the asymmetric decision bounds, which promote early default boundary hits. To test these predictions, we measure the mean response time difference between correct and incorrect trials for each mouse, and test for a significant difference at the group level. This revealed a significant “slow error” effect (p = 0.009), as predicted by the model. We also found that the average difference in response times between trials where mice either do or do not follow their default response is significant at the group-level effect (p = 0.023).

**Figure S4:**
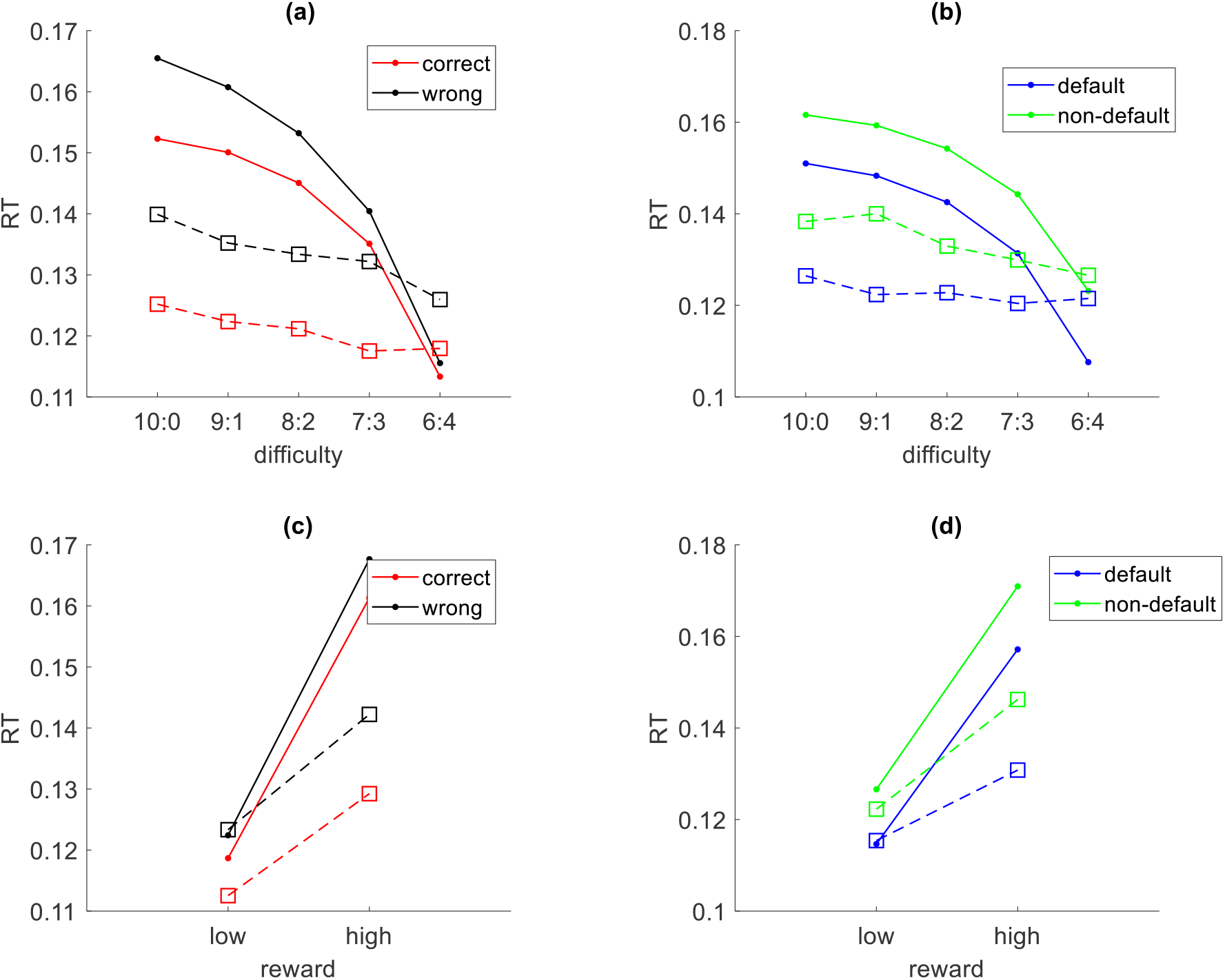
Effect of reward and difficulty on response times. The model predicts opposite motivational modulations of response times as a function of difficulty (panels **a** and **b**) and reward (panels **c** and **d**), irrespective of whether trials are split according to correct/wrong (panels **a** and **c**, red: correct, black: wrong) or default/non-default (panels **b** and **d**, blue: default, green: non-default) responses.

Besides, the decision control model predicts that reducing the reward at stake or increasing decision difficulty renders the cost/benefit balance unfavorable, which eventually shrinks optimal stopping boundaries. This tends to shorten the duration of controlled processing of the visual cue, irrespective of whether responses eventually follow the cue and/or the default bias. To test this prediction, we regressed response times for each mouse against trial difficulty and reward at stake, having split trials according to either correctness (correct versus incorrect response) or default-following (stay versus switch responses). Regression coefficients were then analyzed at the group level. The results of this analysis are summarized on Figure S4. As predicted, we observed a significant positive effect of reward and a significant negative effect of difficulty on overall response times (p < 0.001 and p = 0.005, respectively). Response times for both correct and incorrect trials showed the same pattern, with significant decreases with difficulty (p = 0.018 and p = 0.004) and significant increases with reward (p < 0.001 and p < 0.001). In addition, difficulty significantly reduced response times for non-default responses (p = 0.008), although no significant effect was observed for default responses (p = 0.203). Conversely, reward significantly increased response times for both default and non-default responses (p < 0.001 and p < 0.001, respectively).

## Appendix 5 Variational Bayesian model inversion

For completeness, we performed a variational Bayesian inversion of the model, which served to derive the quantitative model predictions that can be eyeballed on the figures of the Results section. Under our design, reward and difficulty conditions are important because they modify the cost/benefit tradeoff of control regulation. However, they are not expressed in the native units of the model. We thus reparametrize the model to encode reward and difficulty conditions as follows:

- In our design, the reward conditions are encoded through the number of pellets at stake (*p*). Here, we set the reward at stake to be a simple affine mapping of *p*, as follows: *R* = *R*^(0)^ + *R*^(*p*)^ × *p*/max(*p*), where *R*^(0)^ and *R*^(*p*)^ are positive parameters that quantify reward offset and reward sensitivity, respectively.
- Similarly, the difficulty conditions are encoded through the difference in vertical and horizontal lines (Δ*n*). This has two impacts on the model simulations (see Appendix 2). First, the mean evidence signal increases with Δ*n*. Second, the perceptual noise variance *σ*_*s*_ decreases with |Δ*n*|. Accordingly, we simulate the perceptual signal as a series of random samples of a Gaussian variable with the following mean and variance: *E*[*S*] = *s*^(0)^ + *s*^(Δ*n*)^ × Δ*n*/max(|Δ*n*|) and 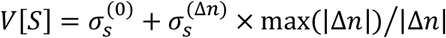. Here, 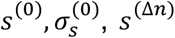 and 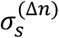 are positive parameters that quantify the signal mean and variance offsets and their sensitivity to difficulty, respectively.

The rest of the model parameters (i.e. *α*_*C*_, *ν*_*c*_, *λ* and 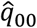) are kept in their native form.

For each of the 10 reward × difficulty conditions, optimal decision bounds were computed, and 2,000 evidence trajectories are simulated as a Bayesian cue-denoising process of the corresponding random perceptual signal samples *S*_*t*_ (where *t* ≤ 200), for both default-congruent (Δ*n* > 0) and default-incongruent cues (Δ*n* < 0). The details of this process are given in the Appendix 2. Each evidence trajectory eventually reaches one of the two bounds: which bound is hit and when it is hit defines the response type (correct/incorrect and default/nondefault) and the response time, respectively. For each, combination of reward, difficulty and default-congruency, response times are binned into deciles, and the within-bin averages of response time and correct/default response rates are quantified. These are the sufficient statistics that are compared to empirical data when adjusting the model parameters using standard variational Bayesian analysis *(Daunizeau et al., 2014)*. In doing so, model inversion relies on the observed quantitative relationship between response time and correct/default response rates. Note that all the model parameters are given unitary prior mean and variance except for *λ* and 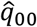, whose prior mean are set to 0.95. The results of this analysis are summarized on Figure S5 below, focusing on the rate of default responses (which correspond to correct responses in default-congruent trials and incorrect responses in default-incongruent trials).

**Figure S5:**
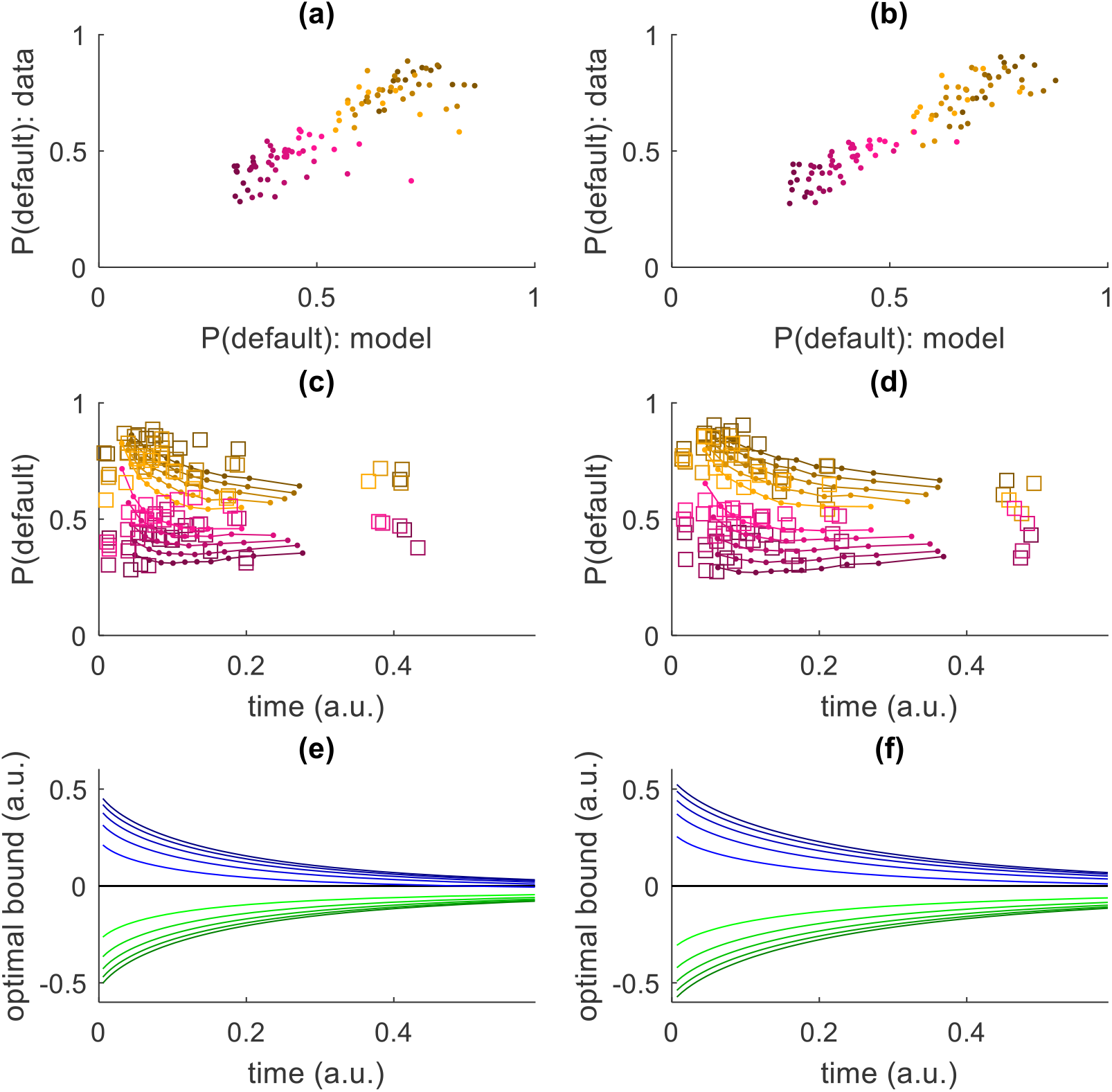
Quantitative data analysis under the optimal control regulation model. Analysis results are shown for each reward condition separately (**a, c, e**: low reward, **b, d, f**: high reward). **(a**,**b)** The observed within-RT-decile average rates of default responses (y-axis) plotted against its model prediction (x-axis). Orange and magenta dots are for default-congruent and default-incongruent trials, respectively, and lighter colors encode task conditions of increasing difficulty. **(c**,**d)** The within-RT-decile average rates of default responses (y-axis, lines: model predictions, square dots: data) plotted against their corresponding average RTs (x-axis). **(e**,**f)** The underlying optimal evidence bounds (x-axis) plotted against decision time (blue: default-compliant bounds, green: default-overriding bounds). Note that the bounds converge at the inferred temporal horizon *H*, which lies beyond the temporal window displayed here.

Despite a slight tendency to compress the distribution of response times, the model explains 80.1% of the variance in within-RT-decile average rates of default responses (Fig S5a and Fig S5b) and 80.5% of the variance in the corresponding average RTs. This level of predictive accuracy is achieved with only eight model parameters, of which just three govern the effects of task conditions. Note that the percentage of explained variance that we report here cannot be directly compared to those reported in Appendix 1 (cf. credit assignment model), because the trial binning in RT-deciles that we use here necessarily increases the effective signal-to-noise ratio of the fitted data when compared to trial-by-trial analyses.

The difference in observed default rates between default-congruent and default-incongruent trials decreases when decision difficulty increases (see Fig S5c and Fig S5d). Under the model, this is due to the dynamics of the underlying optimal evidence bounds, which shrink as decision difficulty increases (Fig S5e and Fig S5f). One can also see that average default response rates increase when decision time decreases (Fig S5c and Fig S5d). Under the model, this is due to the shrinking additional evidence demand for overriding (when compared to confirming) the default response (Fig S5e and Fig S5f). Finally, the (comparatively soft) effect of reward on the default response rate is mostly an indirect consequence of slowing response down (see the slight expansion of evidence bounds between Fig S5e and Fig S5f).

